# Inhibition of Nrf1 activity is a relevant for the HCV-dependent dysregulation of host lipid metabolism

**DOI:** 10.64898/2026.05.29.728748

**Authors:** Olga Szostek, Patrycja Schorsch, Daniela Bender, Eberhard Hildt

## Abstract

**BACKGROUND:** Despite advances in knowledge and medicine, hepatitis C virus (HCV) infection remains a global challenge. The viral life cycle depends heavily on lipid metabolism; consequently, HCV infection is associated with significant alterations to host lipid homeostasis. One of the regulators maintaining this homeostasis is the transcription factor nuclear factor erythroid 2-related factor 1 (Nrf1). There are multiple Nrf1 proteoforms that differ in their capacity to act as a cholesterol sensor or to activate or inhibit gene expression. Previously, we identified that the amount of full-length Nrf1 protein in HCV-replicating cells is significantly reduced. In this study, we investigate whether HCV affects the formation and functionality of the different Nrf1 proteoforms, and whether this contributes to the HCV-dependent dysregulation of cellular lipid metabolism.

**METHODS:** The HCV-Nrf1 crosstalk was characterized using robust methods including: Western blot, FRET Acceptor Photobleaching assay, reverse transcriptase-polymerase chain reaction and confocal laser scanning microscopy-based approaches.

**RESULTS:** HCV infection does not alter the onset of Nrf1 proteoforms generated through proteasomal cleavage of the protein. However, the amount of different Nrf1 proteoforms is significantly reduced in HCV-positive cells due to enhanced Nrf1 turnover. Furthermore, the Nrf1 proteoforms with transcriptional activator functions are prevented from translocation into the nucleus. Reduced Nrf1 activity contributes to elevated cholesterol levels and favors lipid droplets formation. Conversely, rescue of Nrf1 activity in HCV-replicating cells is associated with decreased intracellular cholesterol levels, reduced number of lipid droplets and impaired viral release reflected by intracellular accumulation of the core protein and intact viral particles.

**CONCLUSIONS:** Overall, our results identify Nrf1 as a significant factor in the deregulation of host lipid metabolism induced by HCV. The inhibition of Nrf1 functionality driven by HCV leads to intracellular cholesterol accumulation, resulting in enhanced lipid droplet formation that supports the HCV life cycle and contributes to HCV-associated pathogenesis.

## Background

Despite the efforts undertaken by the World Health Organization (WHO) to eliminate viral hepatitis by 2030, with estimated 50 million of infected individuals, hepatitis C virus (HCV) infection still persists as a global burden [1, 2]. Undeniably, the development of pan-genotypic direct-acting antivirals (DAAs) marked a paradigm shift in the management of the disease. DAAs achieve a sustained virological response (SAV) of over 95% in a short time span of 8-12 weeks of treatment duration [3]. Nonetheless, while the DAAs are highly effective against common epidemic subtypes of HCV present in Europe or North America, their efficacy against rare endemic subtypes prevalent in Africa or Southeast Asia is limited [4]. Moreover, the high price of DAAs restricts their widespread application, particularly in countries with less developed healthcare systems that are burdened by the disease. Another challenging points are HCV underdiagnosis, gaps in knowledge regarding spread of the disease or social and cultural stigma [2, 5]. As a result, globally only 36% individuals with chronic HCV are aware of their status [2]. Being a bloodborne virus, HCV infection spread is mainly common among people who inject drugs (PWID), but it is also connected with unscreened blood transfusions, unsafe medical injections or through unsterilized medical equipment [5–7]. A high proportion of infected individuals (∼70%) fail to clear the virus. In those cases, the acute infection, manifesting with fever, loss of appetite, nausea, dark urine, abdominal pain or jaundice, progresses into chronic infection. This in turn can lead to progressive liver fibrosis, cirrhosis and hepatocellular carcinoma (HCC) [2, 8]. All of the aforementioned factors make the seek for an efficient vaccine against HCV a necessity. However, this necessity is constantly hampered by numerous challenges, starting with the high mutation rate of HCV, genetic variability and quasispecies abundance, viral immune evasion mechanisms or lack of proper animal model [9–11].

Hepatitis C virus is a small enveloped virus with single-strand positive sense RNA as a genome [12]. Its life cycle is strictly confined to the cytoplasm [13]. A hallmark of HCV infection is the formation of “membranous web” (MW), a distinct replication site. MW is primarily derived from the endoplasmic reticulum and is characterized by clustering of single- and double-membrane vesicles (DMVs), viral proteins and RNA as well as host’s cellular factors [14]. The replication sites have unique biochemical properties i.e. are highly enriched in cholesterol, ceramides, phosphatidylcholine, phosphatidylinositol, phosphatidic acid or sphingomyelin [15–17]. Another important organelle for HCV morphogenesis are the cytosolic lipid droplets (cLDs). cLDs are required for the production of infectious viral particles [18]. They act as an assembly platform, connecting the replication and assembly sites of the virus [13, 19, 20]. Moreover, mobilization of lipids from LDs is crucial for viral envelopment and subsequent production of infectious lipo-viro-particles [17]. This being said, HCV induces profound changes in host’s cell lipid metabolism. One of the key mechanisms involved in regulation of intracellular lipid content described by Widenmaier et al. as “guardian of cholesterol homeostasis” is the transcription factor nuclear factor erythroid 2 related factor-1 (Nrf1) [21]. Due to the intimate connection between cholesterol metabolism and the HCV life cycle, this characteristic makes Nrf1 an interesting study subject in the context of HCV infection. The functional interplay of Nrf1 and HCV is so far not characterized. On one hand Nrf1 protects the cells from excess cholesterol by triggering the removal programs, and on the other HCV triggers accumulation of cholesterol within the cell. The Nrf1 transcription factor is a primary member of the Cap’N’Collar (CNC) basic-region leucine zipper (bZIP) transcription factor superfamily [22]. Besides Nrf1, the CNC family comprises of other cytoprotective agents like Nrf2 [23, 24] and Nrf3 [25], p45 NF-E2 [26] and the transcriptional repressors Bach1 and Bach2 [27]. Of note, Nrf2 is heavily restricted by HCV infection. Nrf2 controls the expression of a variety of cytoprotective genes including ones affecting homeostasis of ROS within the cells. Under redox steady conditions Nrf2 is bound to its inhibitor Keap1 and subjected to constant proteolysis. Upon a stress stimuli, namely elevated ROS level, Nrf2 dissociates from Keap1 and is translocated into the nucleus, where it forms a heterodimer with small musculoaponeurotic fibrosarcoma oncogene (sMaf) proteins. The heterodimer binds to the antioxidant response elements (ARE) in the promoter region of cytoprotective genes thereby activating their expression to counteract redox imbalance [28, 29]. However, in HCV-replicating cells an unprecedented mechanism has been observed. Here, the sMaf proteins are withdrawn from the nucleus by the viral NS3 protein. These complexes are bound to cytoplasmatic face of the ER, thereby being trapped in the HCV replication organelles (ROs). Therefore, when Nrf2 is released from Keap1, it is no longer translocated into the nucleus, but instead it binds to the NS3-trapped sMafs in the ROs. Such relocation prevents the Nrf2-mediated regulation of cytoprotective genes [30, 31].

Besides maintaining the cholesterol homeostasis, Nrf1 serves multiple roles in the cells amongst which are: maintaining proteasome homeostasis - since the expression of some proteasomal subunits i.e. PSMB5 depends on ARE elements, induction of anti-inflammatory responses, regulation of embryonic development or defense against reactive oxygen species [32–34]. Such wide functional variety of Nrf1 is intrinsically linked to its structure and subsequent post-translational modifications. The protein is organized into nine structural domains [35] of which the cholesterol recognition amino acid consensus (CRAC) domain is particularly interesting in the context of HCV infection. Through the N-terminally localized CRAC domain, Nrf1 is able to directly bind excess cholesterol, making the protein a cholesterol sensor that promotes cholesterol removal from the cell [21]. This feature however, is preserved for the full-length form of Nrf1. Initially Nrf1 is expressed as ER-bound protein with its C-terminus facing the ER-lumen and the N-terminus protruding into the cytoplasm, typically detected at 140 kDa – 120 kDa. In this state Nrf1 is transcriptionally inactive, yet upon stress stimuli the protein gets activated and undergoes a chain of post-transcriptional modifications [35, 36]. At first the C-terminus is retrotranslocated into the cytoplasm via the p97/VCP complex and subsequently deglycosylated by Peptide-N-Glycanase 1 (NGLY1) [37]. Final step of Nrf1 activation involves a site-specific proteolytic cleavage catalyzed by the aspartyl protease DNA-damage inducible 1 homolog 2 (DDI2) [38]. This step removes the CRAC domain from its structure, releases Nrf1 from the ER and allows its translocation to the nucleus. As a result, Nrf1 gains a variety of proteoforms with different molecular masses ranging from 95 kDa to 25 kDa. Notably, the 85 kDa Nrf1 serves as a main transcriptional activator and the 25 kDa Nrf1 is the dominant-negative proteoform [35, 39, 40]. After translocation to the nucleus, the transcriptionally active Nrf1 proteoforms form heterodimers with sMaf proteins and therefore can effectively recognize the ARE-driven genes and modulate their expression (Fig 1) [36].

**Figure 1.**
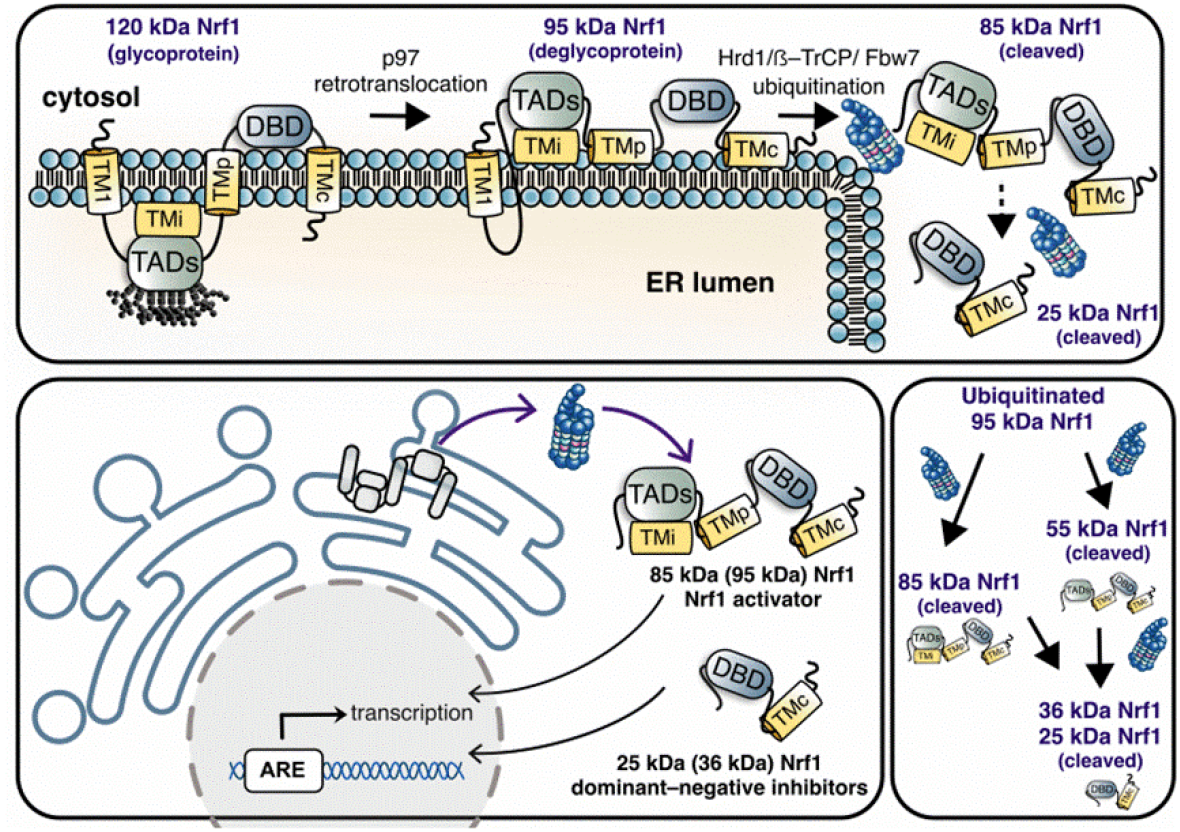
Schematic representation of post-translational modifications of full-length Nrf1. Located in the ER, full-length Nrf1 undergoes various post-translational modification including the proteasomal processing. This allows Nrf1 to be translocated into the nucleus where it mediates the downstream gene expression. During this process Nrf1 gains multiple proteoforms i.e. 95 kDa, 85 kDa, 55 kDa, 36 kDa or 25 kDa.

Our previous work revealed that in HCV-positive cells the amount of Nrf1 protein is significantly lower than in the HCV-negative cells. Moreover, suppression of Nrf1 activity contributes to viral replication, assembly and release of progeny virions. Similarly to Nrf2, the processed and transcriptionally active 85 kDa Nrf1 proteoform can be effectively withdrawn from the nucleus by extracellular sMaf proteins that are trapped in the HCV replication complexes within the ER. This in turn affects the Nrf1-driven expression of ARE-dependent genes, such as NQO1 [41]. Such situation favors the HCV life cycle by maintaining elevated ROS levels within the cells and subsequently contributing to autophagy-dependent release of HCV [42]. HCV morphogenesis critically depends on the virus altering the host’s lipid metabolism. The full-length form of Nrf1 contains the CRAC domain and acts as a cholesterol sensor in cells. Given the significant deregulation of lipid metabolism in HCV-replicating cells, this feature makes Nrf1 a promising target for further study in the context of HCV-induced alterations to lipid metabolism. Therefore, we aim to investigate whether HCV infection impacts the processing pattern of full-length Nrf1, as well as its quantity, as both are crucial for its functionality. Furthermore, we will investigate the implications of HCV-dependent modulation of Nrf1 activity on cellular lipid metabolism and, consequently, on viral morphogenesis. In this context, we will study whether the effect of Nrf1 on viral morphogenesis can be correlated with changes in ‘membranous web’ formation due to deregulated cholesterol/lipid homeostasis.

## Methods

### Cell culture and treatments

For electroporation, highly permissive for HCV infection human hepatoma-derived cell line Huh7.5 (RRID:CVCL_7927) was used [43]. The HCV genomic RNA was synthesized by *in vitro* run-off transcription of linearized plasmid DNA (pFK-pJFH1/GND and pFK-JFH1/J6) using T7 Scribe Standard RNA IVT Kit (Biozym, Germany) according to manufacturer’s protocol. Electroporation with obtained RNA was performed as described elsewhere [44]. Culture media were composed of Dulbecco modified Eagle medium high glucose w/ sodium pyruvate (Bio&Sell GmbH, Germany) supplemented with 2 mM L-glutamine (Bio&Sell GmbH, Germany), 100 μg/mL streptomycin, 100 U/mL penicillin, and 10% v/v fetal bovine serum (Bio&Sell GmbH, Germany). The cells were cultivated at 37°C with 95% relative humidity and 5% CO_2_. HCV-replicating and non-replicating cells were passaged two to three times a week, up to passage seven.

Treatment with 27-hydroxycholesterol (dissolved in 100% v/v ethanol) (MedChemExpress, USA) was performed for 8h at a concentration of 20 µM. The treatment was performed using DMEM supplemented as described above with 0.5% v/v fetal bovine serum (FBS). To sensitize the cells to of 27-hydroxycholesterol and to minimize the uptake of background cholesterol present in FBS, prior to the treatment the cells were starved overnight with DMEM without addition of FBS. Supplementing growth medium with 0.2% v/v ethanol served as treatment for experimental control.

### Plasmids

Plasmids encoding HCV DNA: replication deficient pFK-pJFH1/GND and pFK-JFH1/J6 have been described previously [44–46]. pFK-Luc-Jc1 encodes a bicistronic reporter construct of the full length Jc1 genome [47] and was kindly provided by R. Bartenschlager (University of Heidelberg, Germany). Reporter plasmid pGreenFire1-LXRE harboring the Liver X Receptor (LXR) response elements (LXREs) and the flanking regions in the LXR-α promoter was ordered from IBA Lifesciences. Plasmid encoding EGFP (GenBank: AAB02572.1) was purchased from Clontech Laboratories Inc. Plasmid encoding mCherry (GenBank: AAV52164) was previously generated in our lab. The Δa1pEGFP-N1 (pJo23) vector was purchased from Invitrogen. Plasmid encoding a fusion EGFP-mCherry protein (pcDNA3.1(-)-EGFP-LL-mCherry) as well as plasmid encoding a fusion GFP-mCherry protein with an internal P2A self-cleaving peptide sequence (pcDNA3.1(-)-EGFP-P2A-LL-mCherry) were generated and kindly provided by M. Glitscher (Paul-Ehrlich-Institute, Germany). Three variants of plasmids encoding human full length Nrf1 (endoplasmic reticulum membrane sensor NFE2L1 isoform 2, NP_001317190.1) were purchased from Biomatik Corporation. The first plasmid variant encodes the full length Nrf1 tagged with a C-terminal mCherry (pcDNA3.1 (-) Nrf1-mCherry); second variant encodes full length Nrf1 tagged with a N-terminal EGFP with a signal peptide localizing to the endoplasmic reticulum (cDNA3.1 (-) SignalPeptide-mEGFP-NRF1) and the third plasmid variant encodes full length Nrf1 tagged with a N-terminal EGFP with a signal peptide localizing to the endoplasmic reticulum and a C-terminal mCherry (pcDNA3.1 (-) SignalPeptide-mEGFP-NRF1-mCherry). Plasmid encoding 25kDa Nrf1 C-terminally tagged with EGFP was previously generated in our lab [41].

### Transient transfection

The electroporated Huh7.5 cells were transiently transfected 24h post-seeding using linear polyethyleneimine (PEI) (1 mg/ml) (Polysciences, USA). The cells were seeded at a density of 3×10^5^ cells/well. A DNA:PEI mass ratio of 1:6 was used followed by 16h incubation and medium change. Cells were harvested 48h after transfection, at the peak point of Nrf1 overexpression.

### Luciferase reporter gene assay

To determine the LXR-α promotor activity, HCV-replicating Huh7.5 cells and the control cell were transiently transfected using pGreenFire1-LXRE plasmid (encoding for a LXR promotor-driven firefly luciferase). After 48h post-transfection, the cells were lysed using luciferase lysis buffer (25 mM Tris-HCl, 0.1% Triton-X 100, 2 mM DTT, 2 mM EGTA, 10% glycerol, pH 7.5). The luciferase activity of lysates (50 µl) was analyzed by addition of luciferase substrate (25 mM Tris-HCl pH 7.8, 5 mM MgCl2, 33.3 mM DTT, 0.1 mM EDTA, 470 μM Luciferin, 530 μM ATP) and subsequent detection in the Orion II microplate Luminometer (Titertek Berthold). Technical duplicates of the samples were analyzed. Luciferase levels were referred to the protein concentration of the appropriate lysates, determined by Bradford assay.

### RT-qPCR

The total intracellular RNA was isolated using RNA-Solv® Reagent (Omega Bio-tek, USA) according to the producer’s manual. Equal amount of total RNA (4 µg) was incubated with RQ1 RNase-Free DNase I (Promega, Germany) and reverse transcribed into cDNA using RevertAid H Minus Reverse Transcriptase (Thermo Fisher Scientific, USA). cDNA was thereafter 10-fold diluted and qPCR reaction was performed using the SYBR-green detection kit (Thermo Fisher Scientific, USA) or GreenMasteMix (Genaxxon, Germany). Data were analyzed using the ΔΔCT method and normalized to the amount of RPL27 transcripts.

The following primers were used:

**Table 1.**
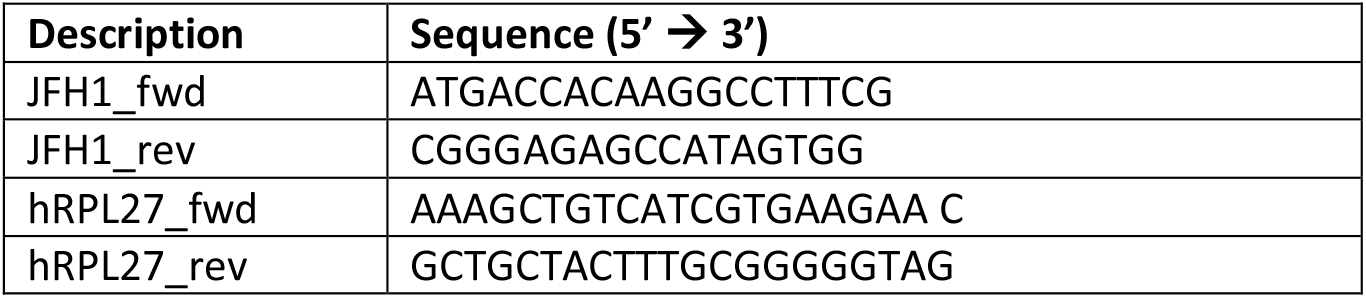
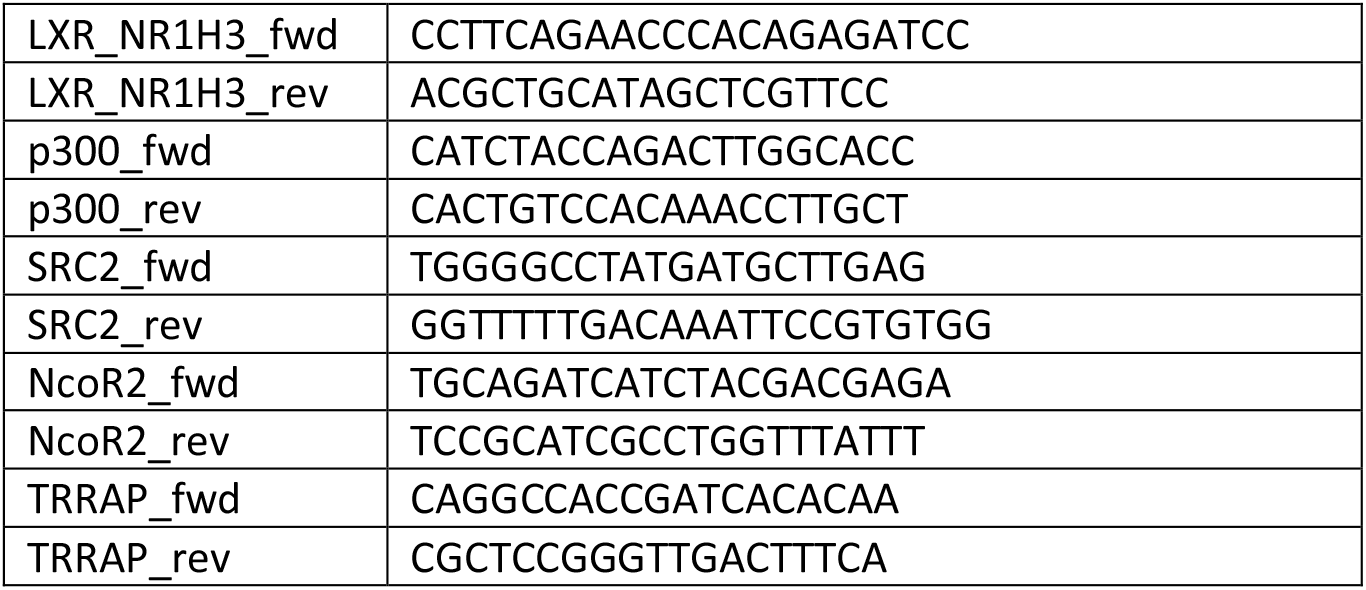
List of primers used for RT-qPCR.

Extracellular viral RNA was isolated from the cell culture supernatants using the QIAmp® Viral RNA Mini Kit (Qiagen, Germany) and detected with LightMix Modular Hepatitis C Virus Kit (TIB Molbiol, Germany) in combination with LightCycler Multiplex RNA Virus Master (Roche Diagnostics, Germany) according to the manufacturer’s protocol. All RT-qPCR experiments were performed using the LightCycler 480 Instrument II (Roche Diagnostics, Germany) and analyzed using the LightCycler480 Software v1.5.1 (Roche Diagnostics, Germany).

### Immunofluorescence microscopy

The cells were grown on glass coverslips (1.5H) (Carl Roth, Germany) and fixed with 4% PFA in PBS at RT for 20 minutes. Next these were blocked with 5% bovine serum (Carl Roth, Germany) in 0.05% TBST for 15 min, unless stated otherwise. Then, the cells were permeabilized with 0.05% TritonX-100 (Sigma-Aldrich, Germany) for 10 min in RT. Primary and secondary antibodies were incubated in humidity chamber for 1h at RT, respectively. For all steps blocking solution was used as a diluent. After washing with 0.05% TBST, the samples were mounted with Mowiol 40-88 (Sigma-Aldrich, Germany). Primary antibodies were raised against HCV-core (MA1-080; Thermo Scientific), NS3 (VIS-1847-100; Biozol/MyBioSource), mCherry (ab167453; Abcam), GFP (632592; Takara). For NS5A detection, polyclonal rabbit-derived serum was used. Secondary antibodies directed against mouse or rabbit were produced in donkey and were conjugated to Alexa Fluor 546 (A10036; Invitrogen or A10040; Thermo Fischer Scientific) or Alexa Fluor 633 (A-21052; Thermo Fischer Scientific). Nuclei were stained with DAPI (4.6-diamidino-2-phenylindole) (Carl Roth, Germany) solution (1 μg/μl) alongside incubation with secondary antibodies. For lipid droplets visualization BODIPY 495/503 solution (Thermo Fisher Scientific, Germany) was added during the incubation with secondary antibodies. Filipin III (Sigma-Aldrich, Germany) was used to stain total intracellular cholesterol. First, the cells were fixed like mentioned above. Afterwards, Schiff-base from FA-fixation was quenched with TBS for 5 min at RT. Cells were then permeabilized as stated above and blocked in 5% bovine serum in TBS containing 0.05 mg/mL filipin. These were then stained with primary and secondary antibodies with addition of 0.05 mg/mL filipin in humidity chamber. After washing with TBS, coverslips were mounted as described above. The stained samples were then analyzed using a Leica Stellaris 8 System (Leica, Germany) using a 93x objective (HC PL APO 93x/ 1.3 Glyc). Images were acquired using LAS X Software (Leica, Germany). All immunofluorescence images were deconvoluted via the LasX Lightning Tool using the Adaptive algorithm. Total fluorescence per cell was calculated using ImageJ software [48] and the following formula: corrected total cell fluorescence (CTCF) = integrated density - (area of selected cell x mean fluorescence of background readings). Unless stated otherwise, a minimum of ten cells were analyzed.

### FRET acceptor-photobleaching

HCV-replicating Huh7.5 cell and the corresponding negative control were transiently transfected as mentioned above. The samples were fixed 48h post-transfection with 4% PFA for 15 min in RT and prepared for immunofluorescence analysis as mentioned before. For HCV detection anti-NS5A specific antiserum was used. FRET efficiency of a single cell was determined using the acceptor photobleaching method on a Leica TCS SP8 confocal microscope with Las X Control Software (Leica, Germany) as described by the manufacturer [49]. Briefly, fixed cells were imaged sequentially for donor (EGFP, excitation 488 nm, emission 500-550 nm) and acceptor (mCherry; excitation 561 nm, emission 580-650 nm) fluorescence before and after selective photobleaching within a defined region of interest (ROI). Acceptor bleaching was performed using the 561 nm laser at high intensity for multiple iterations until >80% photobleaching was achieved. Pixel-wise FRET efficiency was calculated as:

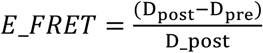

where D_pre and D_post represent EGFP donor intensities before and after mCherry acceptor bleaching, respectively.

### Virus Titration

Virus titers were analyzed based on limited dilution by determining the half-maximal tissue culture infectious dose (TCID_50_) as described elsewhere [50, 51]. For infection analysis, Huh7.5 cells were seeded in 96-well plate at a density of 1x 10^4^ cells/well and infected with serial dilutions of cell culture supernatants, in 6 replicates (5 steps, 1:5 ratio) for 72h. For determination of intracellular TCID_50_, the cells were washed with PBS, trypsinized and pelleted. The pellets were thereafter resuspended in 1 ml of DMEM complete and the cells were lysed by 4-5 freeze/thaw cycles at −80°C and 37°C, respectively. The lysate was centrifuged for 10 min at 17.000xg in the Fresco™ 17 Microcentrifuge (Thermo Fisher Scientific, USA) and the virus containing supernatants were used for infection, as described above. After 72h, cells were fixed with 4% formaldehyde in PBS in RT. Blocking was performed according to the immunofluorescence analysis. HCV-positive cells were detected using NS5A-specific antiserum, overnight at 4°C. Then, the cells were incubated with horseradish peroxidase-coupled donkey-α-rabbit IgG secondary antibody (Healthcare, USA) and stained with 3-amino-9-ethylcarbazol (30mM Na-acetate, 12mM acetic acid, 0.05% w/v 3-amino-9-ethylcarbazol, 0.01% H2O2). The resulting TCID_50_ was calculated according to Spearman and Kärber [52, 53].

### SDS-PAGE and Western blot analysis

For detection of the processing pattern of the full length Nrf1, cell lysates were prepared using RIPA buffer (50 mM Tris-HCl pH 7.2, 150 mM NaCl, 0.1% SDS (w/v), 1% sodium deoxycholate (w/v), 1% Triton X-100) containing protease inhibitors. The lysates were sonicated, equal protein amount (120 µg) was diluted in 1x SDS loading dye and denatured for 10 min in 95°C. The samples were separated by SDS-PAGE electrophoresis containing 8% or 12% v/v acrylamide. Thereafter, proteins were transferred onto polyvinylidene difluoride (PVDF) membrane (Carl Roth, Germany) and blocked in 1x ROTI-Block solution (Carl Roth, Germany). For detection of the endogenous Nrf1 proteoforms, cell lysates were prepared using the SDS-lysis buffer (55.6 mM Tris-HCl pH 6.8, 2.2% SDS (w/v), 11.1% glycerol) containing protease and phosphatase inhibitors. The lysates were sonicated 3 times, diluted in 1x SDS loading dye and denatured for 10 min in 95°C. 50 µl of the sample was subjected to separation by SDS-PAGE electrophoresis containing 8% or 12% v/v acrylamide. After transferring onto PVDF membrane, the unspecific signals were blocked by 5% milk powder solution (Carl Roth, Germany). The chosen proteins were detected by incubation with specific primary and secondary antibodies. All the antibodies were diluted in blocking solution. The following primary antibodies were used: anti-C-terminal Nrf1 (PA590023; Thermo Scientific and sc-28379; Santa Cruz), anti-N-terminal Nrf1 (#8052; Cell Signaling), anti-NS3 (VIS-1847-100; Biozol/MyBioSource), anti-βactin (A5316; Sigma-Aldrich), anti-βtubulin (ABIN7272982; Antikoerper-Online) and anti-mCherry, anti-GFP, anti-NS5A as described above. Secondary antibodies directed against mouse or rabbit were coupled to IRDye680RD or IRDye800CW (926-68073, 926-32213; LI-COR Biosciences) as well as to horseradish peroxidase (NA934, NXA931; GE Healthcare). The specific fluorescence signals were detected using the LI-COR CLx Odyssey Imaging System (LI-COR Biosciences, Germany). After incubation with Luminata Forte Western HRP substrate (Merck Chemicals GmbH Millipore GmbH, Germany), chemiluminescence was detected with INTAS-Imaging System (Intas Pharmaceuticals, India). Densitometric analyses were performed using the LI-COR Image Studio Lite v5.2.5 after normalization to β-actin or β-tubulin.

### Lipid droplet analysis

LDs analysis was performed using particle analysis feature in Fiji (Image J) open source analysis software [48]. Size of the particle was set as 0.01-infinity (inch^2). Circularity was set as 0.00-1.00. The number of lipid droplets for each cell was counted. Minimum of ten cells were analyzed.

### Endoplasmic Reticulum analysis

For ER visualizations primary antibody raised against calreticulin (AB92516-1001; Abcam) was used and labeled with a secondary antibody conjugated to Alexa Fluor 488 (A-21206, Invitrogen). The samples were prepared as stated above. Fixation was performed using methanol to preserve the calreticulin immunostaining. The obtained confocal images were analyzed using Fiji (Image J) open source analysis software [48]. The workflow was adapted from Lu, van Tartwijk et al. [54] and modified accordingly. In brief, the images were pre-processed using the Median filter and background subtraction. Next, the images were imported to Trainable Weka Segmentation [55] to separate ER network and the background. Trained images were binarized and skeletonized by AnalyzeSkeleton Fiji plugin. For every condition, minimum of 4 cells were analyzed. The cell was divided into 4-5 regions of interest (ROIs) of the same size, which were treated technical replicates. Obtained measurements were averaged and the number of ER branches, junctions, end point voxels and triple points alongside with average branch length were analyzed.

### Statistical analysis

Each figure legend shows the number of independent experiments corresponding to the figure. Prism 9.2 software (GraphPad Prism version 9.2 for Windows, GraphPad Software, USA, www.graphpad.com) was used to perform statistical analysis and to plot the graphical representation of the data. Results are described as the mean ± standard error of the mean (SEM). For statistical comparisons, normality test was performed using the Shapiro-Wilk conditions. For normally distributed data statistical significance was calculated by the unpaired t-test. Data not showing Gaussian distribution were tested using the Mann-Whitney test. Statistical significance is displayed as stated: ns—not significant, *p < 0.05, **p < 0.01, ***p < 0.001, ****p < 0.0001. Threshold for the p value was set using the Holm-Šídák method.

## Results

### HCV infection does not alter the cleavage pattern of the full-length Nrf1

Recently we have described, that there is an intense crosstalk between HCV and the Nrf1 transcription factor. We found that HCV infection leads to a significant decrease in the amount of the full-length Nrf1 [41]. Effective induction of ARE-driven gene expression by Nrf1 requires, among other post-translational modifications, proteasomal processing for conversion of the ER-membrane associated full-length Nrf1 into transcriptionally active soluble forms. Thereby, it gains multiple isoforms referred to as proteoforms [56]. Here, we investigate whether HCV infection affects Nrf1 processing, which is important for its functionality. For this purpose, HCV-replicating cells and the control cells were transfected with a construct enabling expression of the full-length Nrf1 C-terminally tagged with mCherry. Since the processing canonically occurs at the N-terminus, the resulting distinct cleaved Nrf1 proteoforms are still C-terminally tagged, which enables their detection by Western blot. We observed no differences in the pattern of generated proteoforms regardless of HCV infection (Fig 2A). In addition, no proteoforms with a molecular mass different from those previously reported in the literature were detected. Yet, we observed that the amount of each proteoform in comparison to HCV-negative cells is strongly reduced (Fig 2A). This reduction in the amount of different Nrf1 proteoforms could be confirmed by comparative analysis of endogenous Nrf1 in HCV-positive *versus* HCV-negative cells (Additional file 1, S1 Fig).

**Figure 2.**
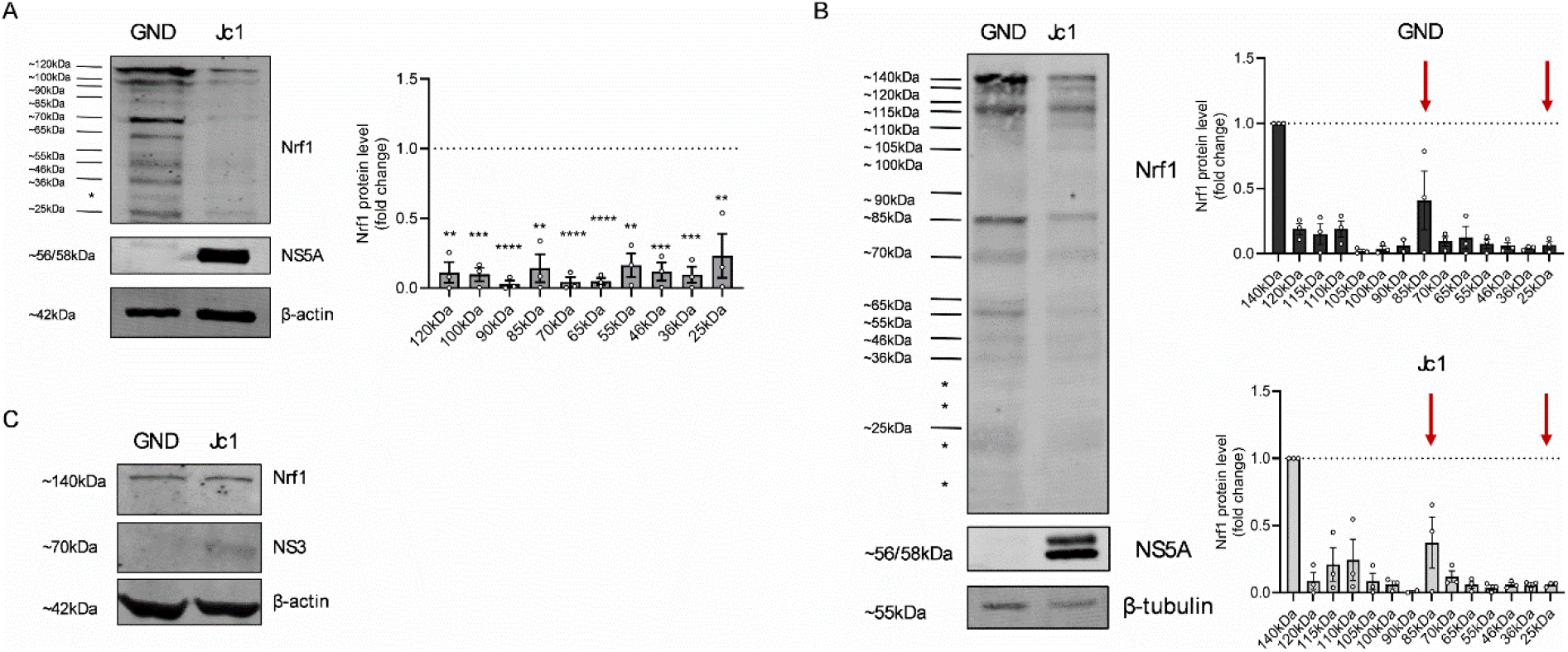
The cleavage pattern of full-length Nrf1 remains unaffected regardless of HCV infection. (A) Representative Western blot and the respective densitometric analyses of cellular lysates derived from stably HCV-replicating (Jc1) cells and the corresponding negative control (GND). For visualization of the cleavage pattern of full-length Nrf1, mCherry specific antibody was used. In addition, NS5A was detected to confirm HCV replication. Detection of β-actin served as a loading control. Unspecific lane indicated by *. Relative values are referred to the control cells (GND) (set to 1); N=3 biological replicates. (B) Representative Western blot and respective densitometric analyses of cellular lysates derived HCV-replicating (Jc1) cells and the corresponding negative control (GND). For visualization of the cleavage pattern of full-length Nrf1, mCherry specific antibody was used. In addition, NS5A was detected to confirm HCV replication. Detection of β-tubulin served as a loading control. Unspecific lane indicated by *. The profile of produced proteoforms in GND (upper) and Jc1 (lower) was calculated referring to the full-length 140 kDa Nrf1 proteoform, set to 1. The active 85 kDa Nrf1 proteoform and the trans-dominant negative 25 kDa Nrf1 proteoform indicated by arrows. N=3 biological replicates. (C) Representative Western blot analysis of cellular lysates derived from stably HCV-replicating (Jc1) cells and the corresponding negative control (GND). Lack of C-terminally cleaved Nrf1 proteoforms was confirmed by visualization of GFP-specific signal. In addition, NS3 was detected to confirm HCV replication. Detection of β-actin served as a loading control. N=3 biological replicates. For all experiments, statistics were performed as mean ± SEM, unpaired t-test referred to control with ns—not significant, *p < 0.05, **p < 0.01, ***p < 0.001, ****p < 0.0001.

Overexpression of the dominant negative 25 kDa Nrf1 fragment in HCV-negative cells, contributed to deregulation of cellular kinome profile and resembled the kinome profile of HCV-positive cells [41]. In light of this, we asked if this is due to an increased formation of the trans-dominant negative proteoforms of Nrf1. For this purpose, we quantified to amount of each produced endogenuous proteoform in HCV-positive and HCV-negative cells, respectively and referred it to the full-length 140 kDa Nrf1 amount. Nonetheless, the analysis did not reveal a shift towards enhanced formation of the trans-dominant negative proteoforms in HCV-positive cells. The most predominantly generated Nrf1 proteoform in both HCV-positive and HCV-negative cells is the 85 kDa Nrf1 (Fig 2B). Next, to deepen the analysis we investigated whether HCV-infection can induce C-terminal cleavage of Nrf1. For this purpose, we transfected the cells with a vector expressing full-length Nrf1 that is tagged with EGFP at the N-terminus. If the processing occurs at the C-terminus of Nrf1 a variety of different proteoforms should be detected on the Western blot using an EGFP-specific antibody. After subjecting the cell lysates to Western blot analysis, we detected exclusively the full-length Nrf1 proteoform (Fig 2C).

The results indicate that HCV infection does not induce changes in the cleavage pattern of full-length Nrf1. Furthermore, Nrf1 processing occurs exclusively at the N-terminus of the protein, regardless of HCV infection. The profile of generated proteoforms remains consistent compared to HCV-negative cells. However, the amount of each Nrf1 proteoform produced in HCV-positive cells is significantly lower than in uninfected cells. These data suggest that the inhibitory phenotype of HCV on Nrf1-dependent gene expression is not due to an increased formation of the 25kDa fragment, but rather to other mechanisms, such as a decreased amount of Nrf1 or the previously described delocalisation of sMaf proteins.

### Enhanced processing of Nrf1 in HCV-positive cells

To investigate whether the decreased amount of Nrf1 in HCV-positive cells is due to an increased turnover of the protein, we decided to perform a FRET acceptor-photobleaching assay. For this purpose, we transfected the cells with a vector expressing the full-length Nrf1 harboring an N-terminal EGFP and C-terminal mCherry, making it suitable for FRET-based experiments. For a precise analysis independent from the transfection efficiency, analysis was performed on a single cell level. Briefly, upon bleaching the mCherry tag serving as acceptor, we expected to note a change in the fluorescence signal of EGFP which served as a donor in our system. Indeed, the experiment revealed that the calculated FRET efficiency was significantly decreased in the HCV-positive cells in comparison to HCV-negative control (Fig 3A). We interpret this result as meaning that HCV infection induces the processing of the active Nrf1 proteoform, which is characterized by the N- and C-termini being oriented towards the cytosol. FRET is only possible if both termini face the same cellular compartment. This is the case for the 95kD form, which is the precursor of the active forms. At first glance, these results appear to contradict our previous observation that the half-life of Nrf1 remains unaffected in HCV-positive cells. However, the half-life of Nrf1 was determined based on full-length Nrf1, represented by the inactive 120kD /140kD Nrf1 forms provides a more detailed interpretation of these results.

**Figure 3.**
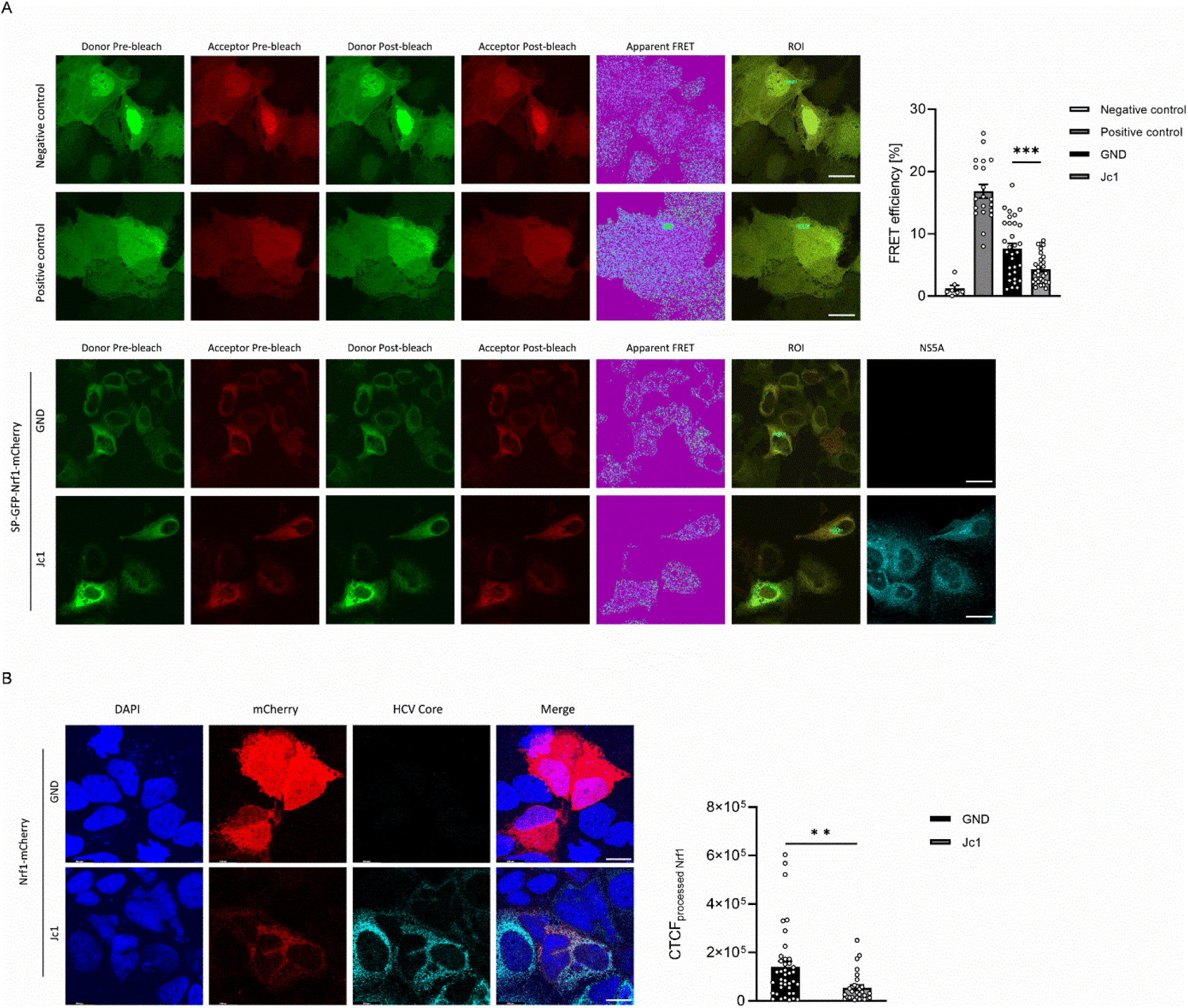
Induced full-length Nrf1 processing in HCV-positive cells. (A) Representative cell images obtained by conventional acceptor photobleaching FRET. Stably HCV-replicating (Jc1) cells and the corresponding negative control cells (GND) were transfected with vector expressing SP-EGFP-Nrf1-mCherry construct, where EGFP is the fluorescence donor and mCherry is the fluorescence acceptor. Additionally, the cells were transfected with a vector expressing EGFP-mCherry as a positive control for FRET and with a vector expressing EGFP-mCherry with an internal P2A self-cleaving peptide sequence as a negative control for FRET. Regions of bleaching indicated by a green box (ROI). NS5A was detected to confirm HCV-replication (cyan). Scale bar, 23.2 µm. FRET efficiencies were calculated with Las X Control Software during mCherry photobleaching. Images are representative of 3 biological replicates, with number of cells analyzed ≥7. (B) CLSM analysis of stably HCV-replicating (Jc1) cells and the corresponding negative control cells (GND). The cells were transfected with a vector expressing Nrf1-mCherry. For visualization of Nrf1, the mCherry-specific signal was detected (red). HCV Core was detected to confirm HCV replication (cyan). Nuclei were stained with DAPI (blue). Scale bar, 15.5 µm. Quantification of the relative CTCF of nuclear mCherry signal corresponding to the processed Nrf1. For each condition, a minimum of 29 cells was analyzed. For all experiments, statistics were performed as mean ± SEM, unpaired t-test referred to control with ns—not significant, *p < 0.05, **p < 0.01, ***p < 0.001, ****p < 0.0001.

To investigate if the increased processing correlates with an elevated level of Nrf1-specific proteoforms in the nucleus, we determined the fluorescence intensity of the Nrf1-mCherry signal localized in the cells’ nuclei, as processing solubilizes the membrane-associated full length Nrf1 and is a prerequisite for nuclear translocation (Fig 3B). Quantitative analysis of the nuclear Nrf1-specific fluorescence intensity revealed, that the nuclear localization of Nrf1 was more occurrent in HCV-negative cells (Fig 3B) as compared to HCV-positive cells. This is in accordance with the fact that, processed Nrf1 fragments (85 kDa Nrf1) can be withdrawn from the nucleus by extracellular sMafs trapped in the replicon complexes, as initially described for Nrf2 [30].

### Overexpression of full-length Nrf1 leads to a decrease in total cholesterol content in HCV-positive cells

HCV life cycle is strictly connected to the host’s cell lipid metabolism, with lipid droplets being a crucial microenvironment for viral morphogenesis. On the contrary, Nrf1 is known to counteract excessive lipid accumulation through triggering lipid removal programs. Moreover, we previously demonstrated that Nrf1 activity is impaired in HCV-positive cells [41]. Taking all these aspects into consideration, we investigated what impact the diminished amount and reduced activity of Nrf1 has on certain aspects of lipid metabolism in HCV-positive cells and what are the consequences for the HCV life cycle.

First, we evaluated the total cholesterol content in HCV-positive and -negative cells that overexpressed full-length Nrf1 in comparison to Mock transfected cells. Our hypothesis stated, that the overexpression of full-length Nrf1 will significantly reduce the total cholesterol content. For this purpose, we overexpressed Nrf1-mCherry fusion protein in HCV-positive and HCV-negative cells. Surprisingly, we did not detect differences in the cholesterol amount regardless of Nrf1 presence. However, we could distinguish three populations of Nrf1 in the cells: (i) with predominant Nrf1 localization in the cytoplasm (ii) in the nucleus (iii) or with equal distribution in both compartments indicating a mixture of processed and unprocessed Nrf1 (Additional file 1, S2 Fig). Correlation of the filipin fluorescence intensity with the localization of Nrf1 revealed that the predominant nuclear localization of Nrf1 correlates with the tendency of an increased intracellular cholesterol level. The nuclear form might represent short Nrf1 proteoforms that lack the sMaf binding domain and act as dominant negative inhibitors. Therefore, the induction of the cholesterol removal program is impaired. If Nrf1-mCherry fluorescence was localized predominantly in the cytoplasm, we noticed a downward tendency in the filipin CTCF in HCV-positive cells in comparison to the Mock control. The cytoplasmic localization might reflect that its sMaf binding domain is still present resulting in a sequestration of Nrf1 to the replicon complexes on the surface of the ER where sMaf is trapped by NS3. A smaller fraction of active Nrf1 due to overload of the sMaf might be able to enter the nucleus and trigger activation of the Nrf1-dependent cholesterol removal program (Additional file1, S2 Fig).

To rule out possible contradictory effects of the transcriptionally active trans-dominant negative proteoforms, we decided to repeat the experiment, this time using a vector expressing Nrf1 tagged with EGFP at the N-terminus (SP-EGFP-Nrf1). This allows the observed green fluorescence to be correlated exclusively with the full-length Nrf1. In HCV-positive cells we observe a significantly higher intracellular level of cholesterol which is reduced by overexpression of Nrf1 (Fig 4). In HCV-negative cells the already much lower intracellular lower cholesterol level is not so prominently further decreased by overexpression of Nrf1 (Fig 4). This suggests that in HCV-negative cells, endogenous Nrf1 maintains intracellular cholesterol levels without the need for overexpression.

**Figure 4.**
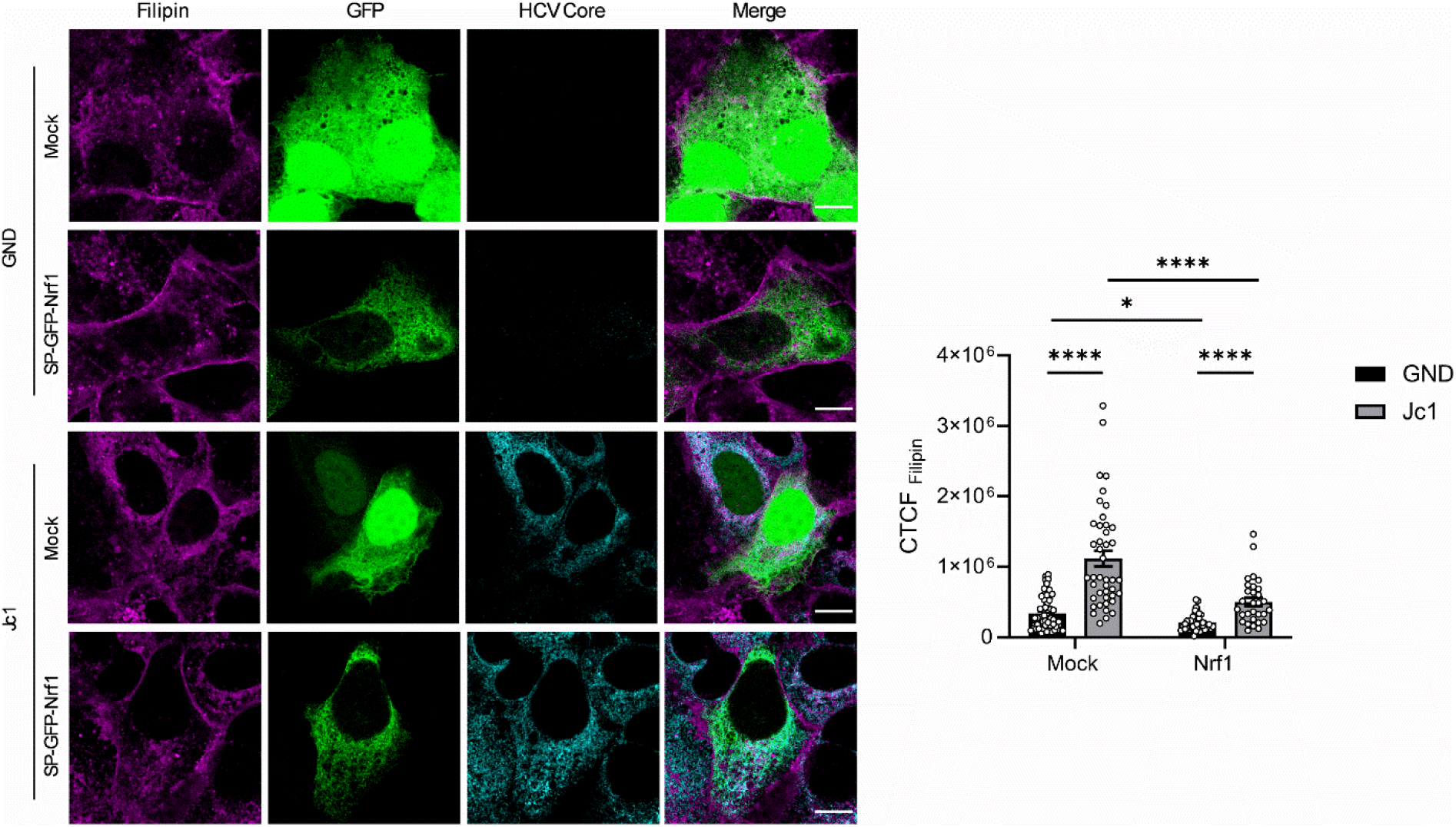
Full-length Nrf1 overexpression significantly decreases intracellular cholesterol content in HCV-replicating cells. CLSM analysis of stably HCV-replicating (Jc1) cells and the corresponding negative control cells (GND). The cells were transfected with the SP-EGFP-Nrf1 construct or EGFP (Mock). For visualization of Nrf1, the EGFP-specific fluorescence was detected (green). Intracellular cholesterol level was visualized with filipin III (magenta). HCV Core was detected to confirm HCV replication (cyan). Scale bar, 12.4 µm. Quantification of the relative CTCF of filipin signal. For each condition, a minimum of 30 cells was analyzed. For all experiments, statistics were performed as mean ± SEM, Mann-Witney test referred to control with ns—not significant, *p < 0.05, **p < 0.01, ***p < 0.001, ****p < 0.0001.

These results indicate, that in HCV-positive cells the impaired Nrf1 activity contributes to elevated cholesterol level.

### Low activity of LXR-α in HCV-replicating cells results from impaired Nrf1 activity but does not involve additional activators or corepressors

The data above indicate that impaired Nrf1 activity in HCV positive cells correlates with increased intracellular cholesterol level. This raises the question about pathways affected by the impaired Nrf1 activity. Liver X receptor alpha (LXR-α) is a nuclear receptor activated by oxysterols in a case of excess cholesterol. It promotes cholesterol efflux from the cells, thereby restricting the negative effects of cholesterol overload on the cell [57, 58]. As observed in our previous work, LXR-α promotor activity is diminished in HCV-positive cells. However, overexpression of the 85 kDa Nrf1, considered as the paradigm of transcriptionally active Nrf1 proteoform, only slightly rescued the phenotype [41]. Thereby the role of Nrf1 in HCV-mediated inhibition of LXR-α and its relevance for the control of intracellular cholesterol level remains to be elucidated.

Since full-length Nrf1 acts as a sensor for cholesterol changes in the cell, we investigated whether full-length Nrf1 can also rescue the LXR-α activity in HCV-positive cells. In accordance to the previous finding, we observed a significantly lower activity of LXR-α promotor in HCV-positive cells (Fig 5A). Notably, after overexpression of the full length Nrf1, LXR-α promotor activity is significantly enhanced, both in HCV-positive (∼2.47-fold of control) and HCV-negative cells (∼2.5-fold of control). Based on this, we conclude that amongst other proteoforms, expression of full-length Nrf1 acts as relevant up-regulator of LXR-α activity. The reduced amount of full-length Nrf1 in HCV-replicating cells can be correlated to the observed significantly weaker LXR-α activity upon HCV infection.

**Figure 5.**
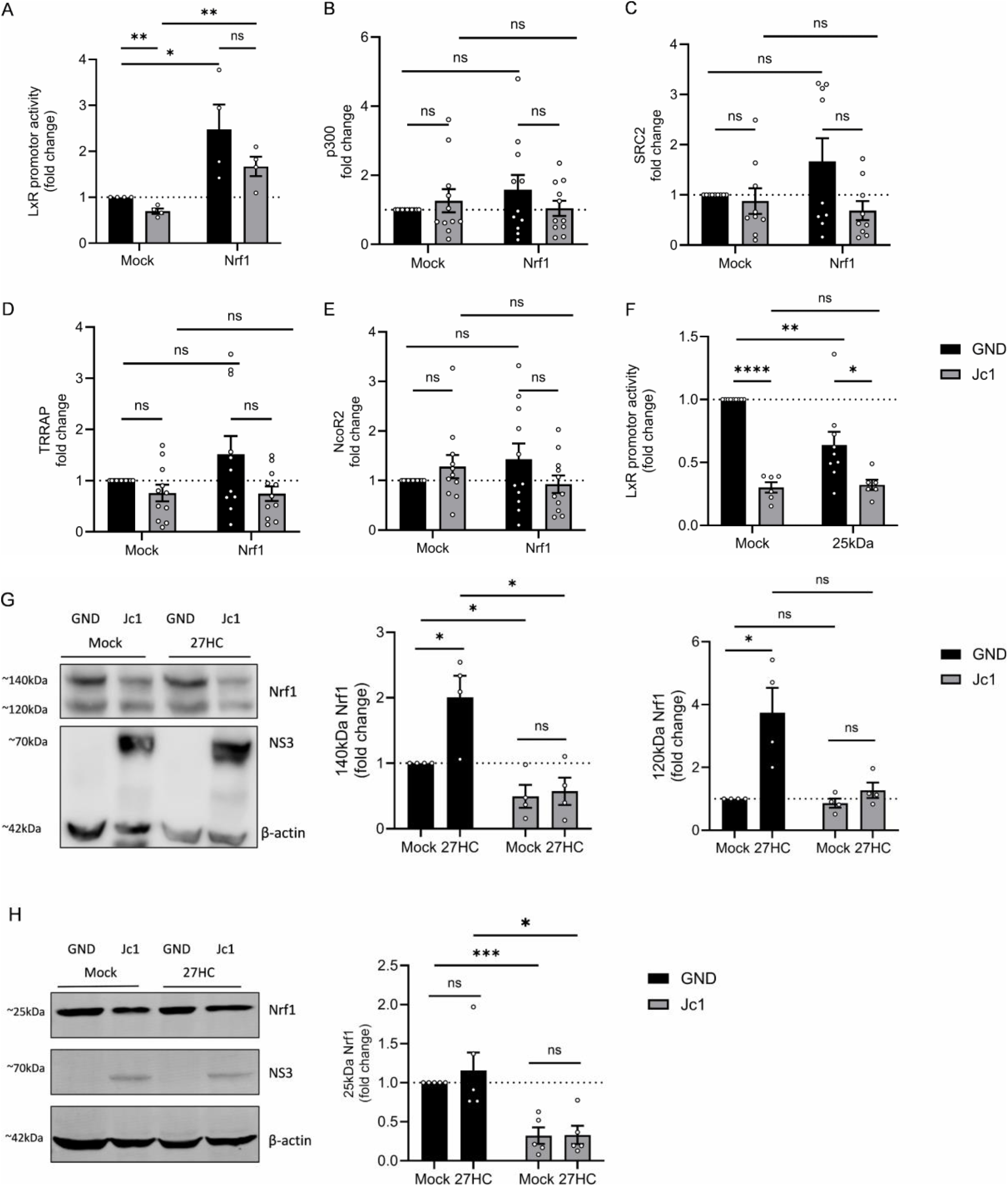
Impaired LXR-α activity results from diminished full-length Nrf1 functionality. Stably HCV-replicating (Jc1) cells or corresponding negative control cells (GND) were transfected with a vector expressing Nrf1-mCherry or witch a vector expressing EGFP-mCherry (Mock). (A) The cells were co-transfected with a reporter construct expressing the luciferase gene under control of the LXR-α promoter. Relative luciferase activity is referred to the Mock transfected cells (set to 1). N=4. (B-E) qPCR analyses to monitor the relative changes in the number of specific transcripts for LXR-α co-activators: p300, SRC2, TRRAP or inhibitor: NcoR2. Relative values are referred to Mock transfected cells (set to 1). N≥9. (F) Stably HCV-replicating (Jc1) cells or corresponding negative control cells (GND) were transfected with a vector expressing Nrf1-25kDa-GFP or Mock transfected (pJo23). The cells were co-transfected with a reporter construct expressing the luciferase gene under control of the LXR-α promoter. Relative luciferase activity is referred to the Mock transfected cells (set to 1). N≥5. (G/H) Representative Western blot and the respective densitometric analyses of cellular lysates derived from stably HCV-replicating (Jc1) cells and the corresponding negative control (GND). The cells were treated with 27-hydroxycholesterol or Mock treated with 0.2% ethanol for 8h. (G) Full-length Nrf1 was detected using an Nrf1-specific antibody binding to the N-terminal part of the protein. N=4. (H) 25 kDa Nrf1 was detected using an Nrf1-specific antibody binding to the C-terminal part of the protein. N=5. In addition, NS3 was detected to confirm HCV-replication. Detection of β-actin served as loading control. For all experiments, statistics were performed as mean ± SEM, multiple t-test referred to control with ns—not significant, *p < 0.05, **p < 0.01, ***p < 0.001, ****p < 0.0001.

In order to deepen the analysis, in the next step we checked whether the Nrf1 dependent regulation of LXR-α promoter involves additional partners, that could be affected during HCV infection. For this purpose, we identified coactivators or inhibitors of LXR-α, that could potentially be affected. Among them were: p300 [59], SRC2 [60] and TRRAP [61], which serve as LXR-α coactivators and NcoR2 serving as a repressor [62]. The level of transcripts of every single gene was quantified by qPCR for HCV-positive and HCV-negative cells that were either Mock transfected or transfected with a full-length Nrf1 expression vector. The analysis revealed, that in HCV-positive cells p300 and NcoR2-specific mRNA levels were slightly increased (Fig 5B, E), while SCR2 and TRRAP mRNA levels were slightly decreased compared to HCV-negative cells (Fig 5C, D). However, the differences between HCV-positive and HCV-negative cells were not significant. Yet, an interesting pattern could be observed: each of the genes is upregulated by overexpression of full-length Nrf1 in HCV-negative cells. This upregulation cannot be observed in HCV-positive cells. On the contrary, in HCV-positive cells expression of the LXR-α inhibitor NcoR2 is slightly diminished after Nrf1 overexpression. However, the effect is not significant.

These data suggest that among the analyzed modulators there is no single co-activator or inhibitor that selectively acts in cooperation with Nrf1 regulating the LXR-α activity.

According to Widenmaier et al. nuclear Nrf1 represses a transcriptional program that otherwise defends against cholesterol accumulation. This program includes LXR-driven cholesterol export pathways. In this model, Nrf1 indirectly affects LXR-α by repression of normally LXR-driven cholesterol export programs, thus loss of nuclear Nrf1 leads to derepression of LXR-α activity and altered cholesterol homeostasis [21].

Based on this model we came up with the following hypothesis: different Nrf1 proteoforms act in certain equilibrium to regulate LXR-α activity and the equilibrium is dysbalanced in HCV-positive cells. Naturally, the full-length form of Nrf1 acts as a sensor for the cholesterol excess in the cell. As proteoform responsible for LXR-α repression we propose 25 kDa Nrf1. In accordance to this, overexpression of 25 kDa Nrf1 proteoform significantly decreases the activity of LXR-α promotor in HCV-negative cells (Fig 5F). This indicates, that under these experimental conditions, the 25kDa Nrf1 fulfills its role as the trans-dominant negative proteoform of Nrf1. However, in HCV-positive cells the decrease could not be observed. This is most likely due to the already prominently decreased promotor activity in the Mock transfected HCV-positive cells (Fig 5F). Further downregulation of the LXR-α promoter activity by the trans-dominant negative Nrf1 proteoform is no longer possible.

To corroborate the hypothesis, we examined the endogenous Nrf1 levels with particular consideration of the full-length Nrf1 proteoforms, as well as 25 kDa Nrf1. For this purpose, the cells were subjected to the cholesterol overload with 27-hydroxycholesterol. The chosen concentration of the treatment was 20µM since upon this treatment conditions, the overall cholesterol content increases significantly, without interference in cell viability (Additional file 1, S3 Fig). Under these conditions a significant increase of 120 kDa and 140 kDa Nrf1 proteoforms in HCV-negative cells was observed by Western blot analysis. In case of HCV-positive cells a minimal increase of these proteoforms was found. This reflects the impaired overall Nrf1 activity, including its cholesterol sensing function in HCV-positive cells. In HCV-negative cells upon stimulation with 27-hydroxycholesterol the induction of the 25 kDa Nrf1 proteoform is much less pronounced as compared to the full-length form. This reflects the functionality of the Nrf1-depenedent cholesterol removal program. In case of the HCV-positive cells there is no significant impact on the level of the 25 kDa proteoform. Again, this might be due to the multiple inhibition of the Nrf1 system in HCV-replicating cell.

Taken together, these data indicate that in HCV-positive cells the cholesterol sensing function of Nrf1 and its role in control of intracellular cholesterol level is impaired.

### Increased lipid droplet amount in HCV-positive cells as a functional implication of impaired Nrf1 activity

Cytosolic lipid droplets (LDs) represent another key aspect of cellular lipid content and play a central role in the HCV life cycle. Under normal conditions they function as a storage organelle engaged in maintaining lipid homeostasis [63]. However, upon HCV infection the lipid homeostasis is disrupted also on the lipid droplet level. HCV infection triggers LDs relocation in the perinuclear region of the cell. Here LDs serve as a crucial microenvironment for viral morphogenesis as they are a bridging platform for the viral replication and assembly sites [12, 64]. Taking this relevance into consideration, in the next step we investigated, whether impaired functionality of full-length Nrf1 influences the number of cytosolic LDs in HCV-positive cells. In HCV-positive cells the number of LDs is increased (average of ∼24 LDs/cell) in comparison to HCV-negative cells (average of ∼10 LDs/cell). Moreover, overexpression of full-length Nrf1 significantly reduces the amount of lipid droplets in the HCV-positive cells (average of ∼6 LDs/cell). The observed reduction was not so prominent in the Nrf1 overexpressing HCV-negative cells (average of ∼6 LDs/cell) (Fig 6).

**Figure 6.**
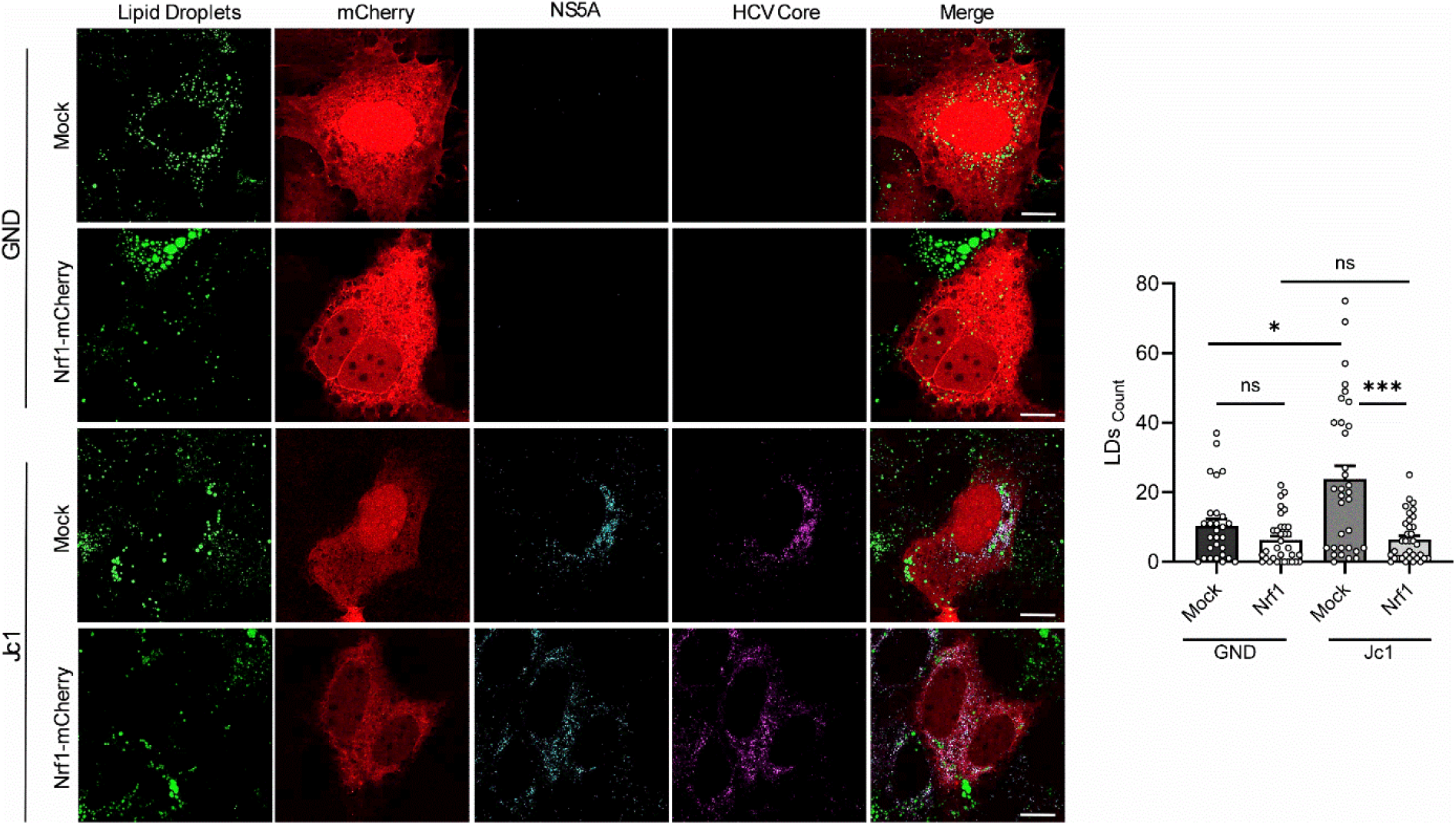
HCV-induced changes in Nfr1 activity affect lipid droplet number. CLSM analysis of stably HCV-replicating (Jc1) cells and the corresponding negative control cells (GND). The cells were transfected with the Nrf1-mCherry construct or mCherry (Mock). For visualization of Nrf1, the mCherry-specific fluorescence was detected (red). Lipid droplets were stained with BODIPY 495/503 (green). NS5A (cyan) and HCV Core (magenta) were detected using specific antisera. Scale bar, 15.5 µm. The number of lipid droplets was determined using Fiji (Image J) software. For each condition, a minimum of 28 cells was analyzed. For all experiments, statistics were performed as mean ± SEM, Mann-Witney test referred to control with ns—not significant, *p < 0.05, **p < 0.01, ***p < 0.001, ****p < 0.0001.

These data once again underline the interplay between impaired Nrf1 functionality, lipid metabolism and LD formation. Rescue of Nrf1 functionality in HCV-positive cells leads to a significant decreased number of LDs. In light of the relevance of LDs for the viral life cycle this suggests that the impaired Nrf1 functionality via this interplay could favor HCV replication.

### Overexpression of full-length Nrf1 negatively impacts HCV release

To further assess the impact of Nrf1 on the HCV life cycle we evaluated the intra- and extracellular viral genomes in the cells overexpressing the full-length Nrf1 proteoforms *versus* the Mock transfected cells. In parallel, the number of intracellular and released viral particles were evaluated in the same settings. Overexpression of the full-length Nrf1 proteoform leads to a significant decrease in both intracellular (∼0.71-fold of control) and extracellular (∼0.77-fold of control) viral genomes (Fig 7A, B). Interestingly, it leads to a significant decrease in the amount of released (∼0.37-fold of control) viral particles, but at the same time the amount of intracellular viral particles raises significantly (∼1.73-fold of control) (Fig 7C, D). Moreover, despite full-length Nrf1 overexpression, viral replication rate stays constant as evidenced by luciferase reporter virus (Fig 7E). This indicates that rescue of full-length Nrf1 functionality primarily affects viral release and not replication.

**Figure 7.**
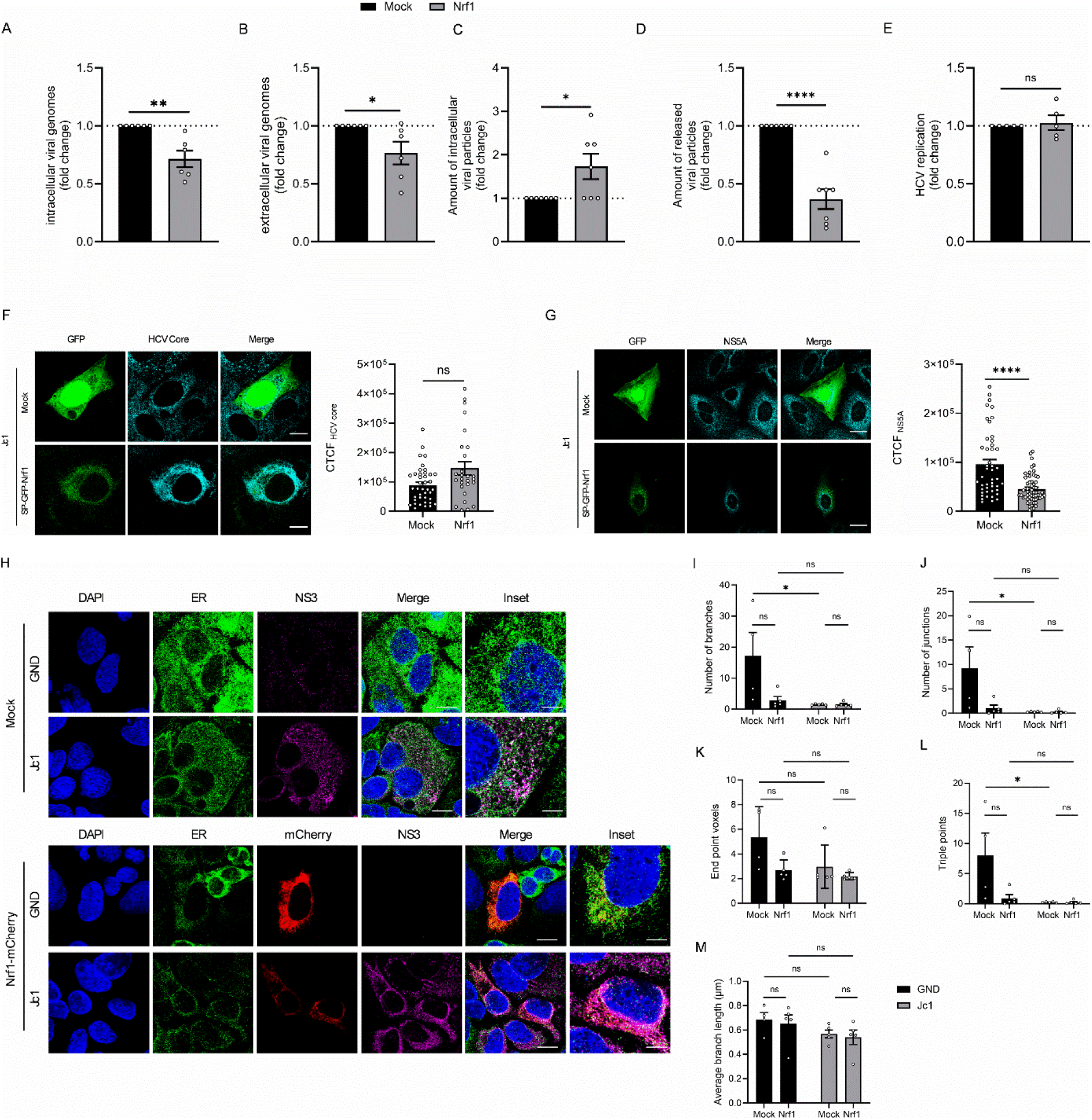
Full-length Nrf1 interferes with HCV release. Stably HCV-replicating (Jc1) cells were transfected with a vector expressing the Nrf1-mCherry construct or Mock transfected with EGFP-mCherry. (A/B) qPCR analyses to monitor (A) intracellular and (B) extracellular viral genomes. Relative values are referred to the Mock transfected cells (set to 1). N=6. Detection of intra-(C) and extracellular (D) viral particles by TCID_50_. Relative values are referred to the Mock transfected cells (set to 1). N=7. (E) Luciferase reporter gene assay of Huh7.5 cells replicating an HCV-luc reporter construct. The cells were transfected with a vector expressing the Nrf1-mCherry construct or a vector expressing EGFP (Mock). Relative values are referred to the Mock transfected cells (set to 1). N=5. (F/G) CLSM analysis of stably HCV-replicating (Jc1) cells. The cells were transfected with the SP-EGFP-Nrf1 construct or EGFP (Mock). For visualization of Nrf1, the EGFP-specific fluorescence was detected (green). (F) HCV Core was detected using specific antibody (cyan). Scale bar, 12.4 µm. Quantification of the relative CTCF of HCV Core. For each condition, a minimum of 28 cells was analyzed. (G) NS5A was detected using specific antibody (cyan). Scale bar, 24.9 µm. Quantification of the relative CTCF of NS5A. For each condition, a minimum of 46 cells was analyzed. (H) CLSM analysis of stably HCV-replicating (Jc1) cells and the corresponding negative control cells (GND). The cells were transfected with the Nrf1-mCherry construct or mCherry (Mock). Due to the fixation with methanol, mCherry fluorescence was abolished. For visualization of Nrf1, the mCherry-specific fluorescence was detected (red). ER was visualized using calreticulin specific antibody (green). NS3 (magenta) was detected using specific antisera. Nuclei were stained with DAPI (blue). Scale bar, 12.4 µm; inset scale bar, 5.5 µm. Number of (I) branches, (J) junctions, (K) end point voxels, (L) triple points as well as (M) average branch length was determined using Fiji (Image J) software. For all experiments, statistics were performed as mean ± SEM, unpaired t-test or Mann-Witney test referred to control with ns—not significant, *p < 0.05, **p < 0.01, ***p < 0.001, ****p < 0.0001.

Based on the results obtained above we questioned, whether through induction of cholesterol removal program the full-length Nrf1 interferes with “membranous web” structure, leading to impairments in viral assembly/release and thereby to accumulation of viral particles intracellularly. To verify this, we first conducted immunofluorescence analysis and quantified the fluorescence intensity of the HCV core in the Nrf1 transfected cells in comparison to the control cells. Based on measured CTCF, we found out that the amount of HCV core was slightly increased in the Nrf1 transfected cells (Fig 7F). Simultaneously, we checked whether the tendency is the same when it comes to non-structural viral protein NS5A. In contrast to HCV core, which is a structural viral protein, NS5A amount was significantly decreased in the cells overexpressing full-length Nrf1 (Fig 7G). To further investigate the impact of Nrf1 on the structures of the MW, we decided to examine the ER structure in HCV-positive *versus* HCV-negative cells upon full-length Nrf1 overexpression. For this purpose we fluorescently labelled calreticulin, which can serve as the ER marker [65] (Fig 7H) and characterized the ER structure by determining the number of branches, junctions, end point voxels (branch ends), triple points/three-way junctions and the average branch length. The data revealed that the ER structure in HCV-positive cells is visibly altered in comparison to HCV-negative cells. The number of branches and junctions (Fig 7I, J) is significantly reduced in HCV-positive cells. Similar reduction is observed in end point voxels and triple points in the ER structure of HCV-positive cells (Fig 7K, L). Moreover, we observed a slight yet non-significant reduction of average branch length in HCV-positive cells in comparison to negative control (Fig 7M). Such changes reflect lower ER integrity which can be correlated to the HCV-induced changes and formation of “membranous web”. Upon Nrf1 overexpression, we observe a reduction in the number of branches, junctions, branch ends as well as three-way junctions in the HCV-negative cells (Fig 7I-L). However, this trend cannot be observed for HCV-positive cells, conversely all of the aforementioned ER parameters stay consistent. No changes were noted in the average branch length upon Nrf1 overexpression neither in HCV-positive nor HCV-negative cells (Fig 7M). This outcome leads to the assumption, that by modulation of cholesterol content, Nrf1 heavily affects the structure of ER in HCV-negative cells. On the contrary, in HCV-positive cells the initial extensive changes in the ER structure cannot be rapidly remodeled upon short term overexpression of Nrf1.

Taken together these data indicate that the robust change in ER structure in HCV-positive cells cannot be rapidly converted by rescue of the Nrf1 functionality. In accordance with the stability of the ER structure/”membranous web”, genome replication is not affected by Nrf1 overexpression. However, there is an impaired release as reflected by the elevated level of intracellular core and intracellular accumulation of viral particles while the number of released viral particles is decreased.

## Discussion

Viruses belonging to *Flaviviridae* family share a similarity in modulation the host lipid metabolism in order to support their life cycle [66]. Hepatitis C virus is one among those. Infection with HCV triggers extensive dysregulation in the host cell lipid homeostasis reflected by formation of replication organelles termed “membranous web” and by increased number of cytosolic lipid droplets. Formation of the “membranous web” structure is tightly associated with a strongly increased level of membrane lipids such as cholesterol or phosphatidylcholine [67]. The lipid droplets are important cellular components known to bridge HCV replication and assembly sites [64]. One of the major components of cLD core are cholesterol esters, once again emphasizing the relevance of this class of lipids for HCV life cycle [68]. Identification of the Nrf1 transcription factor as a sensor that directly binds cholesterol via its N-terminally localized CRAC-domain [21] underlines its potent role in counteracting virus-induced modifications. Indeed, in our previous work we report a significantly lower amount of the full-length form of Nrf1 [41]. The aim of the current study is to further characterise the complex interplay between HCV and lipid metabolism, focusing on the HCV-dependent inhibition of Nrf1 and its relevance to the deregulation of cellular lipid metabolism and, consequently, the viral life cycle.

Since Nrf1 functionality depends on post-translational proteolytic cleavage resulting in various proteoforms, we primarily investigate the on the one hand the impact of HCV infection on Nrf1 processing reflected by the cleavage pattern. Moreover, we analyze the subcellular localization of the proteoforms, which allows to draw conclusions on their functionality. On the other hand, we characterize how modulation of Nrf1 expression interferes with HCV replication and release.

In order to acquire regulatory functions, the full-length Nrf1 proteoform needs to be proteasomally cleaved at the N-terminus [38]. Our analysis of the cleavage pattern upon HCV infection revealed no distinct changes in the onset of generated proteoforms. However, alongside with the full-length form, the amount of shorter Nrf1 proteoforms was strongly reduced. As revealed by FRET-based assay, one possibility for this reduction is the increased turnover of the full-length Nrf1 in HCV-positive cells. This however, seem not to align with our previous observation that the half-life of Nrf1 is not affected by HCV-infection [41]. A possible explanation of these seemingly contradictory data could lay in the onset of post-translational modifications prior to proteasomal processing. Nrf1 can adopt various membrane topologies within the ER, with transactivation domains (TADs) facing the lumen or the cytoplasmatic side of ER membrane [69, 70]. The FRET-relevant topology is represented by a fraction characterized by the simultaneous cytosolic orientation of N-terminal and C-terminal part of the protein enabling FRET. Moreover, one factor contributing for membrane topology is the glycosylation status of the proteins [71]. As a matter of fact, inactive 120 kDa Nrf1 is a glycoprotein which TADs face the lumen of the ER. In order to be activated, the TADs are relocated to face the cytosolic part of the ER, followed by deglycosylation as schematically depicted in Fig 1. The changes in the topology of N-terminal part of Nrf1 are a determinant of the overall downstream topology of the protein. This leads to generation of the 95 kDa proteoform, with both N- and C-terminus facing the cytoplasm [69, 70]. This altered membrane topology of 95 kDa Nrf1 can be causative for the observed FRET signal and subjected to rapid cleavage. This is supported by the consistent detection of a ∼90 kDa band in HCV-negative cells, detectable both in Nrf1-overexpression experiments as well as in the endogenous settings (Fig 1A; Additional file 1, S1 Fig). On the other hand, the presence of 120 kDa Nrf1 counterbalances the processing and contributes to the equal Nrf1 half-life in HCV-positive and HCV-negative cells. As the fraction of Nrf1 with a cytosolic orientation of the N- and C-termini is relevant to Nrf1 functionality, and FRET is only possible if both termini have the same orientation, we conclude that our FRET data indicate that the half-life of functional, active Nrf1 is affected by HCV.

HCV infection leads to accumulation of the cholesterol intracellularly [12, 16, 17]. In light of this and the relevant role of Nrf1 for regulation of intracellular cholesterol we investigated, whether the reduced amount of Nrf1 and thereby its diminished activity is a factor contributing to the elevated amount of intracellular cholesterol in HCV-replicating cells. Indeed, restoration of full-length Nrf1 activity in HCV-positive cells, by simple overexpression significantly reduced the overall cholesterol content. Moreover, similar effect was observed for the cytoplasmic LDs. Overexpression of Nrf1 contributed to a decrease in the number of LDs in HCV-positive cells. This effect however, was not so prominent in the absence of HCV. In HCV-negative cells the mechanisms regulating cholesterol homeostasis are in place, resulting in the mere decrease of intracellular cholesterol level by Nrf1 overexpression. These results clearly demonstrate, that impaired Nrf1 activity in HCV-positive cells due to decreased number of functional Nrf1 proteoforms and their sequestration on the ER surface because of sMafs delocalization [30, 31, 41] is one of the factors contributing to elevated lipid content in the cells and therefore to HCV pathogenesis.

It has been shown that signaling pathways regulated by LXR-α limit replication of various viruses such as human immunodeficiency virus (HIV) [72, 73], Newcastle disease virus (NDV) [74] or human cytomegalovirus (HMCV) [75]. Moreover, LXR-α agonists GW3965, T0901317 or 24(S),25-epoxycholesterol inhibit HCV infectivity by restricting viral entry [76]. In HCV-positive cells LXR-α promoter activity is downregulated, which favors HCV life cycle [41]. This work aimed to further decipher the impact of Nrf1 on LXR-α activity and the underlying mechanisms. Indeed, the promoter activity can be strongly recovered by full-length Nrf1 overexpression in HCV-positive cells. This once again indicates, that that multiple inhibition of Nfr1 in HCV-positive cells triggers a cascade of complex interactions resulting in the rise in intracellular cholesterol content.

Viral replication organelles, so called “membranous web” are originating from the endoplasmic reticulum. They are highly, up to ∼9-fold, enriched in cholesterol [15]. Under normal physiological conditions, ER is mainly composed of phosphatidylcholine and phosphatidylethanolamine and it contains relatively low amount of cholesterol [77]. Therefore, HCV evolved an adaptive mechanism that allows accumulation of unesterified cholesterol in the NS5A enriched sites [16]. This is possible due to the interaction of phosphatidylinositol 4-kinase III α (PI4KIIIα) with NS5A and NS5B which result in local rise of phosphatidylinositol 4-phosphate (PI4P) levels [78–80]. This in turn triggers the oxysterol-binding protein 1 (OSBP), to deliver cholesterol to the PI4P rich sites [81]. Here, we investigated whether the rescue of Nrf1 functionality, resulting in a decrease in intracellular cholesterol levels, interfere with HCV morphogenesis. Indeed, after overexpression of Nrf1 we observed an intracellular accumulation of infectious viral particles. However, the number of released particles as well as intra- and extracellular viral genomes was significantly reduced. Moreover, we observed no change in HCV replication rate despite the rescue of Nrf1 activity. In accordance to the unchanged HCV replication rate, analysis of the ER structure reflected by the number of branches, junctions, branch ends, three-way junctions and the average branch length revealed that in HCV-positive cells, no massive changes are detectable after rescuing Nrf1 activity. This might reflect stability of the “membranous web” structure which is not rapidly affected by short term changes in the intracellular cholesterol levels due to temporary changes in Nrf1 functionality/amount and thus results in an almost unaffected viral replication rate. This further suggests, that factors other than a structural change of the ER that contribute to the impaired release and intracellular accumulation of viral particles. Similar observations, namely no impact on viral replication or particle assembly yet retention of the infectious viral particles within the cytoplasm, was observed upon inhibition of multivesicular bodies (MVBs) functionality by U18666 which blocks intracellular cholesterol trafficking. Here, the viral particles were accumulated within the exosomal structures [82]. In addition, we observed a reduced amount of viral NS5A protein. NS5A is a key factor that has the ability to bind viral RNA [83]. In complex with Tip47, NS5A targets viral RNA to the surface of lipid droplets, where it could be assembled with HCV core particles [84]. Based on this it is tempting to speculate, that due to the decreased amount of NS5A, *de novo* synthesized HCV genomes cannot be efficiently transferred to the lipid droplets, whose amount is also reduced upon Nrf1 overexpression. Thus, a fraction of the unbound viral genomes is subjected to degradation. The remaining pole of genomes is still high enough to be efficiently assembled into functional viral particles. The accumulation of infectious viral particles can be further connected to impairment in exosome formation which can mediate the release of HCV [82] and thus leads to the intracellular accumulation of assembled particles and simultaneous decrease in the amount of released viral particles.

One limitation of the study is the choice of the hepatoma cell line for the experiments. Huh7.5 cells were chosen because they are highly susceptible to HCV infection and easy to handle in experiments. Indeed, confirming the results obtained in other cell types, such as primary hepatocytes (PHHs), would further strengthen the observations. However, there are several reasons why PHHs were not included in the experimental setup. Firstly, primary hepatocytes are considered a hard-to-transfect cell type [85], which would be a major concern in the context of this study, as it is partly based on transfection experiments. Furthermore, as they are derived from a donor patient, PHHs are highly variable in terms of gene expression and metabolic processes [86]. Additionally, cultivation difficulties, poor permissiveness for HCV infection and the potential for rapid dedifferentiation would further impact the reproducibility of the results. While PHHs closely resemble the in vivo environment of HCV infection, using them in this initial study would introduce too many variables. As this study provides a novel insight into HCV-induced and Nrf1-dependent deregulation of cellular lipid metabolism, we selected Huh7.5 cells as a more controlled initial in vitro model.

## Conclusions

In summary, our study further characterises the complex, multi-level interplay between HCV and lipid metabolism, and the relevance of the Nrf1 transcription factor in this process. Reduced Nrf1 levels and the subsequent decrease in cholesterol sensing activity undoubtedly contribute to the intracellular accumulation of lipids observed in HCV-infected cells. This, in turn, favours HCV morphogenesis, as evidenced by the produced viral progeny. Thus, our data describe Nrf1 as a relevant factor affecting the HCV life cycle, based on its function as a sensor and modulator of intracellular cholesterol levels.

## Abbreviations

ARE: Antioxidant response element
bZIP: Basic-region leucine zipper
cLDs: Cytosolic lipid droplets
CNC: Cap’N’Collar
CRAC: Cholesterol recognition amino acid consensus (domain)
CTCF: Corrected total cell fluorescence
DAAs: Direct-acting antivirals
DMEM: Dulbecco modified Eagle medium
DMVs: Double-membrane vesicles
DNA: Deoxyribonucleic acid
DTT: Dithiotreitol
EGTA: Ethylene glycol-bis(β-aminoethyl ether)-N,N,N′,N′-tetraacetic acid
ER: Endoplasmic reticulum
FBS: Fetal bovine serum
FRET: Förster Resonance Energy Transfer
HCC: Hepatocellular carcinoma
HCV: Hepatitis C virus
HIV: Human immunodeficiency virus
HMCV: Human cytomegalovirus
kDa: Kilodalton
Keap1: Kelch-like ECH-associated protein 1
LXR-α: Liver X receptor alpha
MW: Membranous web
NDV: Newcastle disease virus
NQO1: NAD(P)H quinone dehydrogenase 1
Nrf1: Nuclear factor erythroid 2 related factor-1
Nrf2: Nuclear factor erythroid 2 related factor-2
OSBP: Oxysterol-binding protein 1
PBS: Phosphate-buffered saline
PFA: Paraformaldehyde
PI4KIIIα: Phosphatidylinositol 4-kinase III α
PI4P: Phosphatidylinositol 4-phosphate
PVDF: Polyvinylidene difluoride
PWID: People who inject drugs
RNA: Ribonucleic acid
ROI: Region of interest
ROs: Replication organelles
ROS: Reactive oxygen species
RT: Room temperature
RT-qPCR: Reverse transcription-quantitative polymerase chain reaction
SEM: Standard error of the mean
sMaf: Small musculoaponeurotic fibrosarcoma oncogene protein
SVR: Sustained virological response
TADs: Transactivation domains
TCID_50_: Half-maximal tissue culture infectious dose
WHO: World Health Organization
27HC: 27-hydroxycholesterol

## Ethics approval and consent to participate

Not applicable.

## Consent for publication

Not applicable.

## Availability of data and materials

The datasets used and/or analyzed during the current study are available from the corresponding author on reasonable request.

## Competing interests

The authors declare that they have no competing interests.

## Funding

The study was supported by a doctoral scholarship from the Studienstiftung des deutschen Volkes obtained by Olga Szostek. The funders had no role in study design, data collection and analysis, decision to publish, or preparation of the manuscript.

## Authors contributions

**Conceptualization:** OS, EH.

**Formal analysis:** OS.

**Funding acquisition:** EH.

**Investigation:** OS, PS.

**Methodology:** OS, EH.

**Project administration:** DB, EH.

**Resources:** OS, EH.

**Supervision:** OS, EH.

**Visualization:** OS.

**Writing – original draft:** OS

**Writing – review & editing:** DB, EH.

All authors read and approved the final manuscript.

## Acknowledgements

The authors thank Gert Carra for the excellent technical support. Further they thank Keerthihan Thiyagarajah, Jan Raupach an Kim-Anna Mentchen for the topic-related discussions.

## Preprint Server

The manuscript is posted to BioRxiv preprint server, **doi:** https://doi.org/10.64898/2026.05.29.728748

## Additional files

Additional file 1

- **S1 Fig. HCV infection does not influence endogenous full-length Nrf1 cleavage pattern.** Representative Western blot and the respective densitometric analyses of cellular lysates derived from stably HCV-replicating (Jc1) cells and the corresponding negative control cells (GND). For Nrf1 detection an antibody binding specifically to the C-terminal part of Nrf1 was used. In addition, NS3 was detected to confirm HCV-replication. Detection of β-actin served as a loading control. Relative values are referred to the control cells (GND) (set to 1); N=4 biological replicates.
- **S2 Fig. Nrf1 proteoforms exhibit different effect on total cholesterol content.** CLSM analysis of stably HCV-replicating (Jc1) cells. The cells were transfected with the Nrf1-mCherry construct or mCherry (Mock). For visualization of Nrf1, the mCherry-specific fluorescence was detected (red). Three populations of cells with different predominant Nrf1 localization can be distinguished: nuclear, cytoplasmatic or nuclear and cytoplasmatic. Intracellular cholesterol level was visualized with filipin III (magenta). HCV Core was detected to confirm HCV replication (cyan). Scale bar, 12.4 µm. Quantification of the relative CTCF of filipin signal. For each condition, a minimum of 3 cells was analyzed. For all experiments, statistics were performed as mean ± SEM, t-test referred to control with ns—not significant, *p < 0.05, **p < 0.01, ***p < 0.001, ****p < 0.0001.
- **S3 Fig. 27-hydorxycholesterol concentration and toxicity assessment.** Huh7.5 cells were incubated with different 27-hydroxycholesterol concentrations, varying from 5 µM to 50 µM for 24h. (A) CLSM analysis. Intracellular cholesterol level was visualized with filipin III (magenta). Scale bar, 25 µm. Quantification of the relative CTCF of filipin signal. For each condition, 20 cells were analyzed. (B) Cell viability assessed via a PrestoBlue assay; values expressed as % metabolic activity referred to the experimental control. For all experiments, statistics were performed as mean ± SEM, t-test referred to control with ns—not significant, *p < 0.05, **p < 0.01, ***p < 0.001, ****p < 0.0001.

## Notes

### Competing Interest Statement

The authors have declared no competing interest.

### Summary of Updates

The focus of the ms has been changed to make the relevance of Nrf1 as a novel factor significsntly affecting the HCV-dependent deregulation of lipid metabolism. Moreover the FRET experiments have been explained in more detail.

