## Supplementary material for "Inhibition of Nrf1 activity is a relevant for the HCV-dependent dysregulation of host lipid metabolism": suppl data

**Additional file 1**

Supplementary Figures

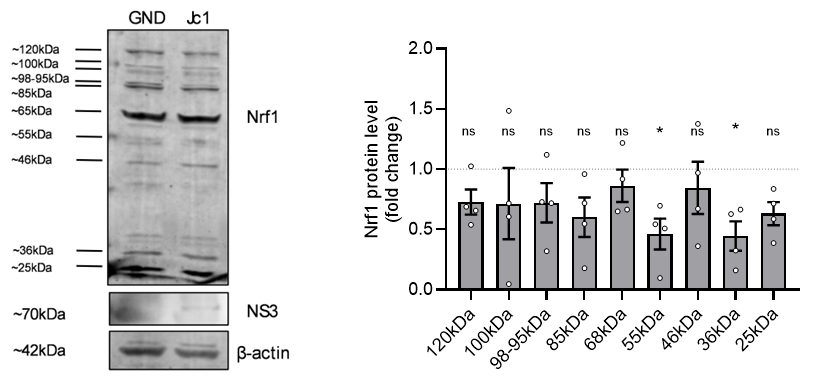

**S1 Fig. HCV infection does not influence endogenous full-length Nrf1 cleavage pattern.** Representative Western blot and the respective densitometric analyses of cellular lysates derived from stably HCV-replicating (Jc1) cells and the corresponding negative control cells (GND). For Nrf1 detection an antibody binding specifically to the C-terminal part of Nrf1 was used. In addition, NS3 was detected to confirm HCV-replication. Detection of β-actin served as a loading control. Relative values are referred to the control cells (GND) (set to 1); N=4 biological replicates.

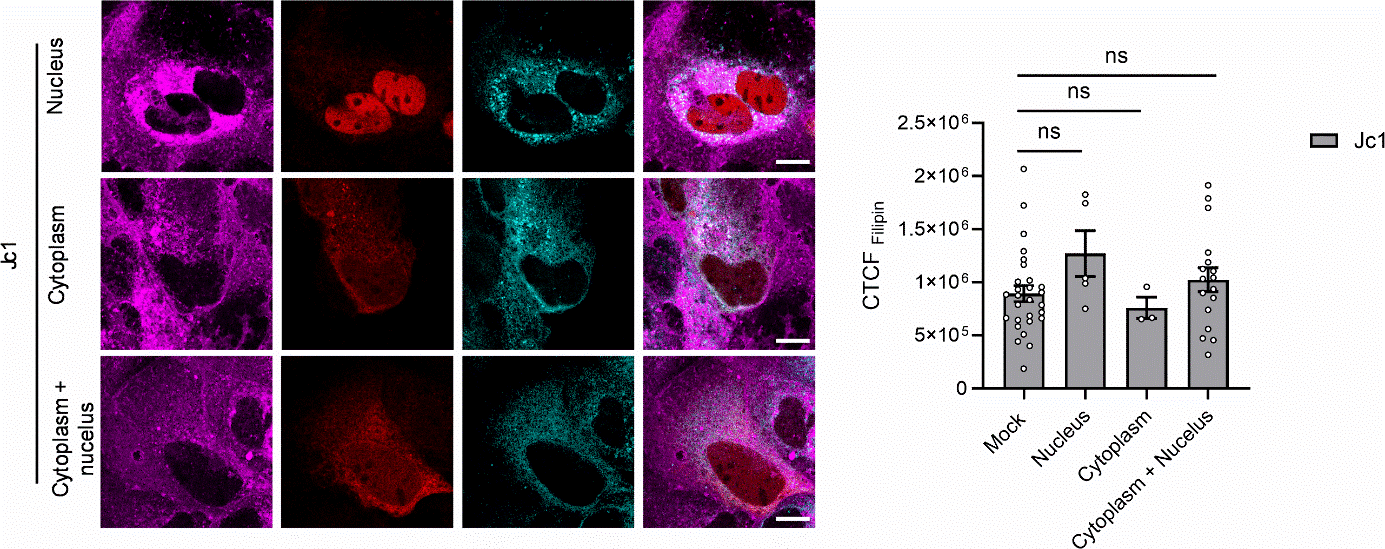
 **S2 Fig. Nrf1 proteoforms exhibit different effect on total cholesterol content.** CLSM analysis of stably HCV-replicating (Jc1) cells. The cells were transfected with the Nrf1-mCherry construct or mCherry (Mock). For visualization of Nrf1, the mCherry-specific fluorescence was detected (red). Three populations of cells with different predominant Nrf1 localization can be distinguished: nuclear, cytoplasmatic or nuclear and cytoplasmatic. Intracellular cholesterol level was visualized with filipin III (magenta). HCV Core was detected to confirm HCV replication (cyan). Scale bar, 12.4 µm. Quantification of the relative CTCF of filipin signal. For each condition, a minimum of 3 cells was analyzed. For all experiments, statistics were performed as mean ± SEM, t-test referred to control with ns—not significant, *p < 0.05, **p < 0.01, ***p < 0.001, ****p < 0.0001.

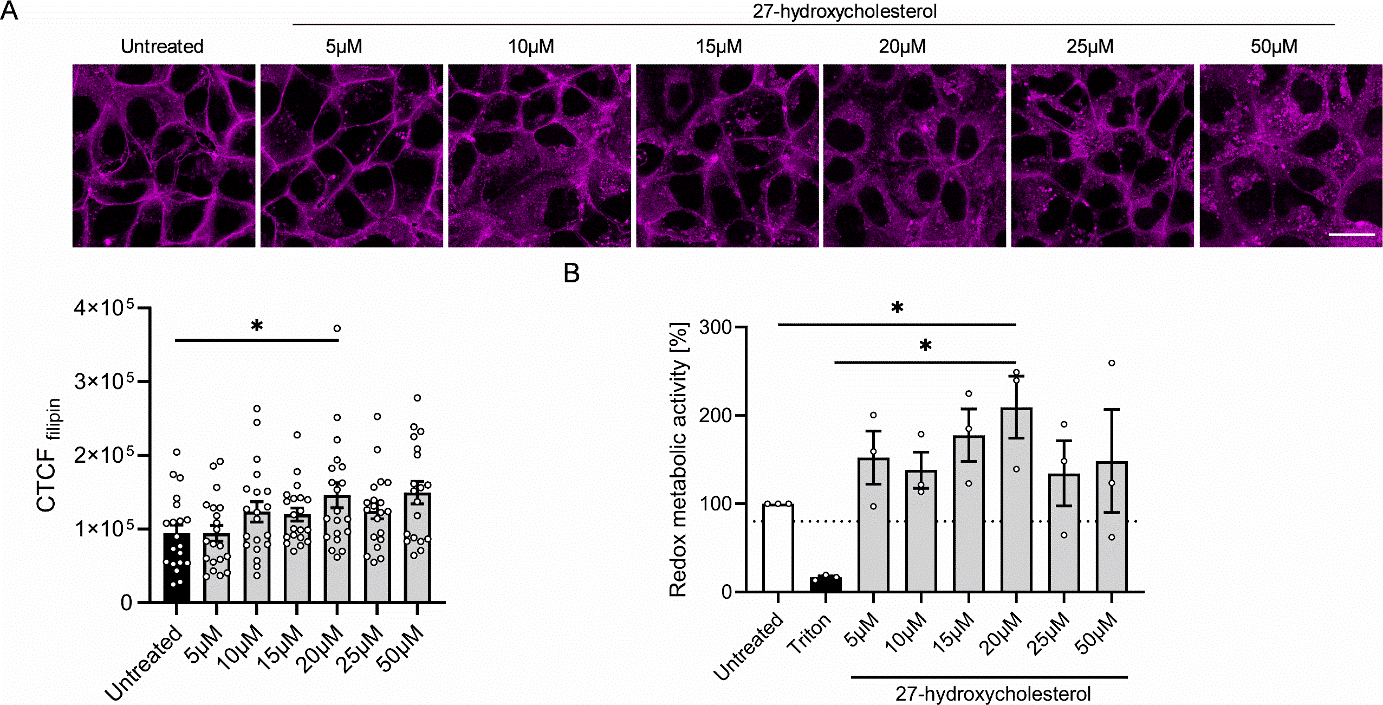

**S3 Fig. 27-hydorxycholesterol concentration and toxicity assessment.** Huh7.5 cells were incubated with different 27-hydroxycholesterol concentrations, varying from 5 µM to 50 µM for 24h. (A) CLSM analysis. Intracellular cholesterol level was visualized with filipin III (magenta). Scale bar, 25 µm. Quantification of the relative CTCF of filipin signal. For each condition, 20 cells were analyzed. (B) Cell viability assessed via a PrestoBlue assay; values expressed as % metabolic activity referred to the experimental control. For all experiments, statistics were performed as mean ± SEM, t-test referred to control with ns—not significant, *p < 0.05, **p < 0.01, ***p < 0.001, ****p < 0.0001.

Supplementary Material and Methods

PrestoBlue assay

The cells were grown in 96‐well‐plate in technical triplicates. Subsequently, the cells were treated with 27-hydorxycholesterol at final concentrations of 5, 10, 15, 20, 25 and 50 µM. Supplementing growth medium with a final concentration of 1 % v/v Triton X‐100 served as qualitative, internal control causing all present cells to disintegrate, therefore leading to the lowest signal possible. Determination of redox‐metabolic activity in living cells was performed using the PrestoBlue™ Cell Viability Reagent (Thermo‐Scientific, Germany) according to the manufacturer’s protocol. Fluorescence of NADH/H+‐dependently formed Resorufin was measured with Spark® Multimode Microplate Reader (TECAN, Switzerland) at Ex_560_/Em_590_. Percentages of redox metabolic activity in treated cells referred to the experimental control were calculated according to Equation below:

$$\% metabolic activity=\frac{\mathrm{RFU}_{treatment}}{\mathrm{RFU}_{control}}\times100$$

RFU = relative fluorescence units
